# Comprehensive benchmarking of somatic single-nucleotide variant and indel detection at ultra-low allele fractions using short- and long-read data

**DOI:** 10.1101/2025.10.13.681545

**Authors:** Yoo-Jin Jiny Ha, Dominika Maziec, Julia Markowski, Stephanie J. Georges, Nancy L. Parmalee, Michele Berselli, Tim H.H. Coorens, Shihua Dong, Stephanie Gardiner, Divya Kalra, Daofeng Li, Benpeng Miao, Rajeeva Musunuri, Liying Xue, Zhi Yu, Kimberly Walker, Lisa Anderson, Natalie Y.T. Au, Carrie Cibulskis, Harsha Doddapaneni, Christopher M. Grochowski, Dana M. Jensen, Tina Lindsay, Kelsey Loy, Azeet Narayan, Giuseppe Narzisi, Jeffrey Ou, Meranda M. Pham, Alexi M. Runnels, Andrew B. Stergachis, Lila M. Sutherlin, Ting Wang, Hu Jin, William C. Feng, Yuwei Zhang, Alexander D. Veit, Clara TaeHee Kim, Hye-Jung E. Chun, SMaHT Network Single Nucleotide Variant (SNV) Working Group, Kristin Ardlie, Robert S. Fulton, Soren Germer, Richard Gibbs, Gabor T. Marth, James T. Bennett, Peter J. Park

## Abstract

Mosaic mutations in normal tissues occur at low variant allele fractions (VAFs), complicating detection. To benchmark strategies, the SMaHT Network created a cell-line mixture (1:49) and produced ultra-deep whole-genome sequencing using short and long reads (five centers, 180–500× each). We assembled a reference of 44,008 mosaic SNVs and 2,059 Indels, cross-validation between platforms to expose limits of short-read analysis. We also partitioned the genome by mappability to examine the impact of genomic context, added a negative reference set, and accounted for culture-derived mutations. When seven institutions applied eleven algorithms to mixture data, call sets were largely discordant across tools and replicates, partly reflecting stochastic presence of low-VAF mutations in biological replicants. For >2% VAF SNVs, sensitivity and precision approached ∼80% at ≥300×, with little gain from additional sequencing. This work provides a comprehensive framework for reliable detection of low-VAF mutations in non-cancer tissues and a valuable resource for the community.

## Introduction

Post-zygotic somatic mutations arise throughout life and are inherited by a subset of cells within an individual, leading to somatic mosaicism^1^. The clonality of somatic mutations varies widely, depending on the timing of acquisition, tissue lineage, and clonal expansion dynamics^2^. Standard bulk whole-genome sequencing (WGS) at ∼30× coverage lacks the sensitivity to detect mutations with low variant allele fractions (VAFs) below 10%, where most mosaic variants fall into, necessitating higher depth of sequencing to capture both early clonal and recently acquired subclonal events. When accurately identified, these mosaic mutations can provide insights into human development, aging, and tissue-specific cellular dynamics. In addition, identification of mosaic variants in non-cancer samples is taking on increasing clinical significance, given the increasing number of pathogenic conditions caused by mosaic mutations in heart^3^, liver^4^, and brain^5^.

However, detecting low-VAF mosaic variants—e.g., those below 5%—remains technically challenging^6,7^ with validation rate remaining modest (∼10-40%). In the presence of sequencing errors and systematic artifacts, the signal-to-noise ratio worsens as VAF decreases, making variant identification progressively more difficult. For cancer mutations, clonal expansion results in increased VAFs, and the availability of matched non-cancer controls enables the filtering of many technical artifacts (e.g., alignment errors shared between tumor and normal samples) as well as shared germline mutations. In contrast, identifying mosaic variants from a single sample without a matched control, as is frequently done in normal tissues, is substantially less accurate due to the inability to distinguish true variants from technical noise and germline mutations^6,7^. The challenge of identifying low-VAF mosaic variants is increasingly important in clinical diagnostics, given the growing number of non-neoplastic conditions such as vascular malformations, overgrowth disorders driven by these types of mutations^8^.

Comprehensive benchmarking of mosaic mutation detection remains limited, especially across the whole genome and those in extremely low VAFs (<5%). Prior studies using cell-line mixtures have yielded valuable insights but were largely confined to detecting relatively high VAF variants in limited genomic regions, e.g., exome or “high-confidence” regions that exclude repetitive or low-complexity sequences entirely^7,9^.

Here, we designed a comprehensive benchmark with the human melanoma cell line COLO829 in an extremely low ratio (tumor:normal ratio of 1:49) and generated a ground reference set of mosaic single-nucleotide variants (mSNVs) and Indels (mIndels). An accurate reference set is made possible by ultra-deep whole-genome sequencing generated by the Somatic Mosaicism across Human Tissues (SMaHT) Network, and ∼300× of PacBio data allowed us to validate the mutations identified from Illumina data, such as those in “low-confidence” regions in which accurate read alignment is error-prone. This orthogonal validation approach substantially increases the quality of the reference set, revealing limitations of Illumina-only benchmarking studies and enabling a comprehensive, genome-wide evaluation of detection methods.

Using the reference set with ultra-low VAFs, we analyze Illumina WGS data sequenced at multiple centers, with a combined coverage of >1,800×. We assessed the consistency of variant calls across several widely used algorithms and evaluated the performance of analysis pipelines from several independent groups within the SMaHT Network. We also discovered the presence of cell culture-induced mutations that are often overlooked in cell line mixture studies. Furthermore, for additional insights on the factors that impact the detection of low-VAF variants, we analyzed a mixture of six well-characterized HapMap samples, for which known germline variants served as mosaic variants to be detected. The cell line mixtures and validated reference sets are publicly available (https://data.smaht.org/) to support the community in developing and benchmarking low-VAF variant detection approaches.

## Results

### Construction of a high-confidence reference set using short- and long-read platforms for a cell line mixture

We created COLO829BLT50, a biologically controlled cell line mixture composed of 2% COLO829 (melanoma) and 98% COLO829BL (matched normal). This design preserved the original germline haplotypes while introducing COLO829-specific mutations at 1% or 2%, corresponding to heterozygous and homozygous COLO829-specific mutations, respectively (**Fig. 1a**). COLO829BLT50 was independently sequenced at five genome sequencing centers (GCCs), with depths ranging from 180× to 500× (**Table S1**), providing a robust and realistic resource for benchmarking low-VAF variant detection.

**Figure 1.**
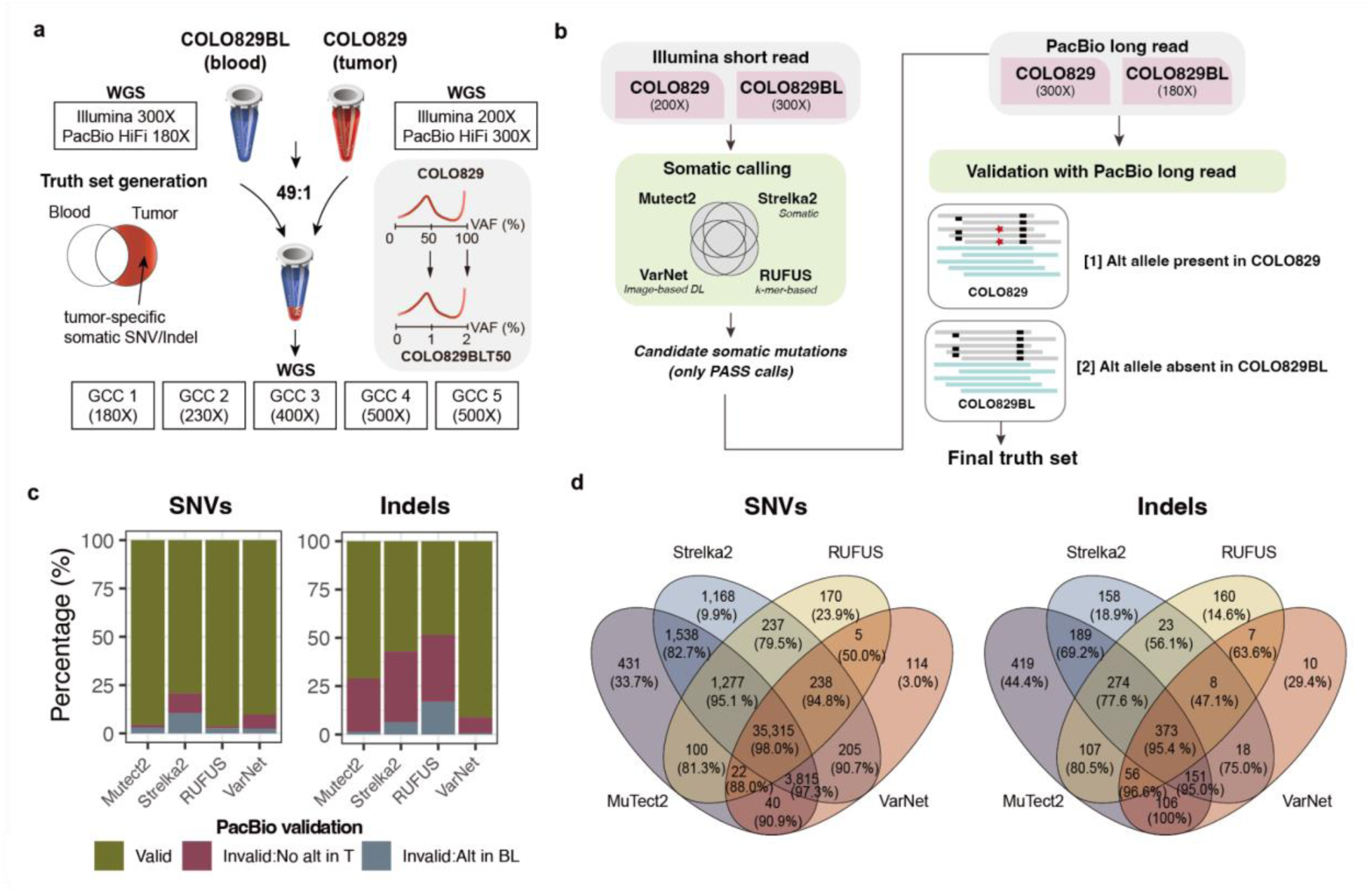
Generation of a high-confidence low-VAF somatic mutation reference set using a melanoma cell line mixture. **a,** COLO829 (tumor) and its matched normal COLO829BL (blood) were mixed in a 1:49 ratio to create COLO829BLT50, a reference material with mosaic mutations at expected VAFs of ∼1–2% and sequenced independently across five Genome Sequencing Centers (GCCs) at 180– 500× coverage. **b,** Reference set generation involves two steps: candidate variant identification from Illumina short-read data using four complementary somatic callers (Mutect2, Strelka2, RUFUS, and VarNet), followed by validation with PacBio long reads. A candidate variant was considered valid if the alternate allele was present in COLO829 and absent in COLO829BL. **c,** PacBio validation outcomes of SNV and indel candidates from each caller. **d,** Venn diagrams showing the overlap among validated SNV (left) and indel (right) calls across the four methods.

To construct the reference set, we first aimed to identify tumor-specific mutations present in COLO829 and absent in COLO829BL. We sequenced each cell line to high depth using Illumina short reads (COLO829: 200×; COLO829BL: 300×) and PacBio long reads (COLO829: 300×; COLO829BL: 180×). A series of quality-control measures were applied to the sequencing data, and key platform-specific features were systematically assessed (**Supplementary Notes** and **Fig.S1**). The reference mutation set generation proceeded in two stages: initial candidate detection using Illumina data, followed by validation with PacBio long reads (**Fig. 1b**).

To maximize sensitivity in initial discovery with short-reads, we applied four complementary Illumina-based methods spanning diverse detection principles: Mutect2 and Strelka2, two widely used somatic variant callers (Mutect2 employs a Bayesian somatic genotyper that uses local assembly of haplotypes, whereas Strelka2 combines a probabilistic somatic model with adaptive indel error estimation and haplotype modeling); RUFUS, a reference-free k-mer-based detector; and VarNet, a deep learning-based image classifier. Variants identified by any of these methods were merged into a unified candidate set, comprising 61,649 SNVs and 4,475 indels.

Although increased sequencing depth improves the quality of variant calls, platform-specific errors persist^10^, especially in genomic regions in which read alignment is imperfect or copy number alterations cause a rare germline variant to appear as somatic^11^. Thus, each candidate COLO829-specific mutation was subsequently evaluated for support in long-read data to eliminate short-read-specific artifacts and other systematic errors. A variant was considered validated if the alternate allele was detected in COLO829 and absent in COLO829BL in PacBio data. Because PacBio reads have a higher per-base error than Illumina and our PacBio data sequenced with high sequencing depth, single read support was too lenient and caused many false positives to be ‘validated.’ Thus, we required a minimum of two supporting PacBio reads. In addition, because long-read sequencing can generate stutter-associated Indel artifacts^12^, particularly near homopolymers and simple repeats, we employed COLO829BL Illumina data to remove variants lacking tumor specificity for Indels.

Given the expected clonality of true tumor-specific mutations in the COLO829 cell line, most SNV candidates were consistently validated using PacBio long reads, confirming their authenticity where short-read-specific artifacts such as misalignment errors were eliminated. Validation rates for SNV were high across methods (median of 92%; range: 79-96%; **Fig. 1c** and **Table S2**). Among the unvalidated calls, a median of 4.3% lacked alternate allele support in PacBio COLO829, while 2.8% were excluded due to the presence of the alternative allele in PacBio COLO829BL.

In contrast, only 63% of Indel candidates were validated (median; range: 48%-91%), with most unconfirmed Indels not supported by tumor long-read data (median: 30.5%). Conversely, a subset of indels-including those identified by methods such as Strelka2 and RUFUS were also found in the matched normal (median: 6.4% and 16%, respectively), suggesting decreased specificity in some genomic contexts. These likely reflect misalignment issues or shared background signals in regions that are more difficult to resolve, highlighting the ongoing challenge of accurate somatic Indel detection in such regions using short-read data alone—even in clonal cell line models.

Validated SNVs and indels from all methods were then combined into the final reference set, comprising 44,005 SNVs, 961 insertions, and 1,098 deletions (**Fig. 1d**). The vast majority of validated SNVs (over 99.5%) were identified by two or more callers out of four, reflecting strong consensus among tools for SNV detection, although some of the variants called by a single algorithm were also validated (e.g., 1,168 were found only by Strelka2). Unlike SNVs, about 36% (747 out of 2,059) of the validated Indels were unique to a single method, underscoring the method-specific sensitivities and the inherent challenges of Indel detection in short-read data. These findings highlight the importance of incorporating diverse variant calling strategies and long-read data to comprehensively capture true somatic variants-particularly Indels-across the genome, enabling both high sensitivity and precision, facilitating rigorous benchmarking of extremely low-VAF mosaic mutations in the COLO829BLT50 model.

### Sampling variability for low-VAF mosaic mutation across independent datasets

To benchmark ultra-low-VAF variant detection performance, we utilized five independent Illumina WGS datasets of the COLO829BLT50 mixture generated at five sequencing centers (GCCs), with depths ranging from 180× to 500× (**Fig. 1a**). We confirmed that the reference SNVs and Indels showed VAFs limited to extremely low levels, mostly under 2%, confirming their origin from mostly copy-number neutral regions in the original cell lines (**Fig. 2a**). For SNVs represented at lower depths (e.g., 180×), VAFs were more tightly concentrated around 1–2%, yielding pronounced peaks that likely reflect limited allele support. With increasing coverage, these peaks were attenuated, and the distributions smoothed. Unlike SNVs, indel VAFs remained widely dispersed even in low-depth datasets, consistent with the increased mapping uncertainty of short reads containing insertions or deletions.

**Figure 2.**
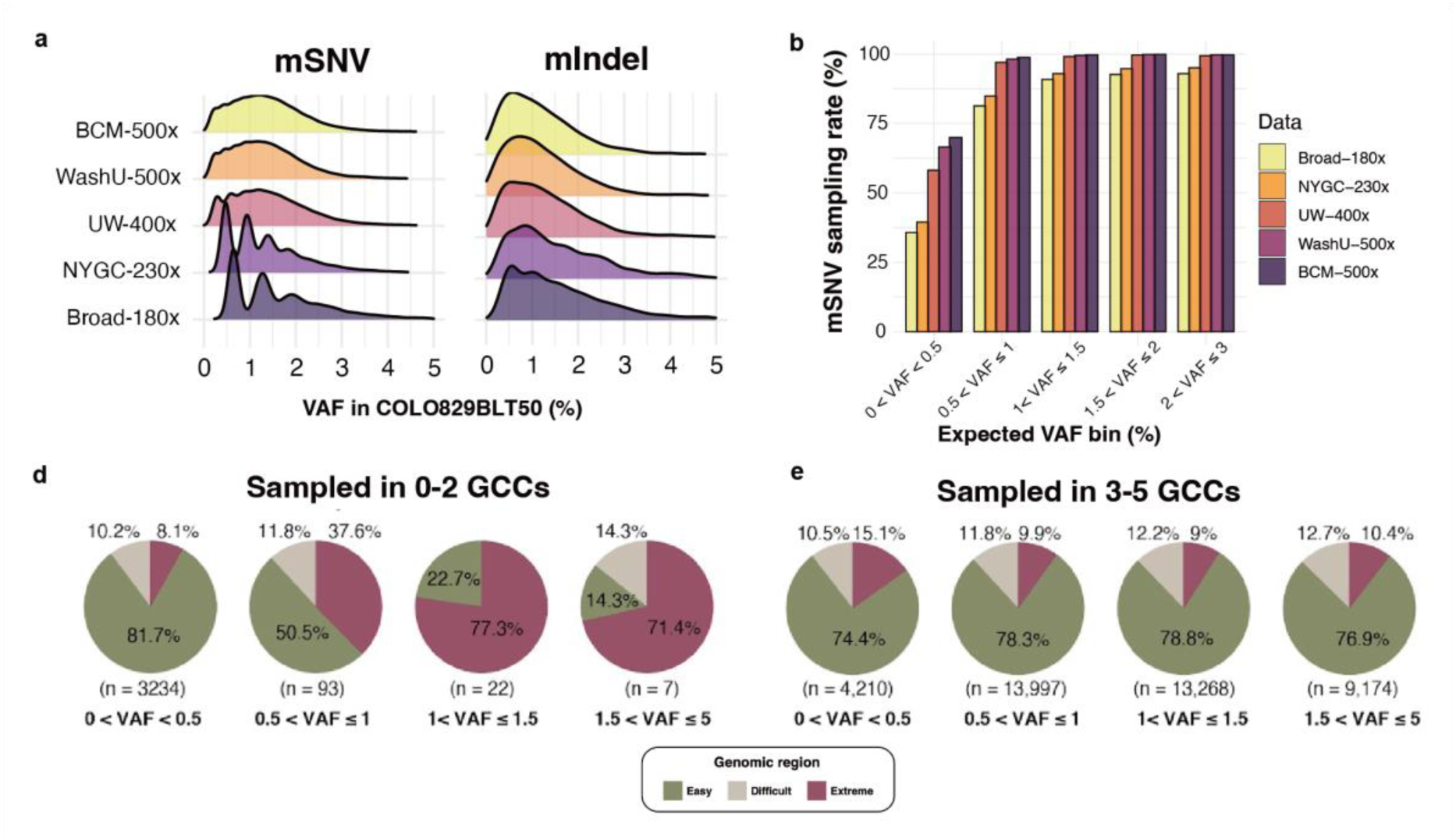
VAF distribution and sampling consistency of reference set variants across independent COLO829BLT50 datasets. **a**, VAF distributions of reference set SNVs and indels as represented in five independently generated Illumina datasets from different sequencing centers (Broad, NYGC, UW-SCRI, WashU-VAI, BCM). Each density plot reflects the observed VAF distribution of reference set variants captured in the respective dataset. **b**, Sampling rate of mosaic SNVs in each dataset, stratified by expected VAF bin. **c**, Variants sampled in ≤2 datasets. **d**, Variants sampled in ≥3 datasets. Pie chart labels indicate the proportion of variants per region and total variant count (n).

Characterization of low-VAF mosaic mutations in bulk sequencing data poses two major challenges: (i) the mutation must be “sampled” during sequencing such that the alternate allele is represented in the data, and (ii) the variant caller must correctly identify the signal and retain it through filtering artifacts. To evaluate the first challenge-allele sampling, we assessed how many reference SNVs and Indels were actually observed in each of the five datasets generated from independent sequencing experiments. As expected, the low-VAF SNV sampling rate was highly dependent on the VAF of each variant (**Fig. 2b**). At 180× depth, >82% of SNVs with VAF >1% were sampled, whereas datasets at 500× achieved near-complete (∼99%) sampling.

However, for variants with VAF <0.5%, even 500× data captured only ∼70% of them, and datasets at 180–230× captured fewer than 40%. These findings highlight that ultra-low-VAF subclonal mutations are intrinsically difficult to capture in bulk sequencing, as variant fragments may be missed even at very high coverage. Sequencing depth must therefore be optimized to balance sensitivity and cost, with biological replicates offering greater benefit than further increases in depth.

We next examined whether genomic context further shapes the sampling of low-VAF variants. Using a stratification framework that partitions the genome into “easy” (74%), “difficult” (10%), and “extreme” (16%) regions^11^, we identified genomic context as a second major determinant of sampling consistency, alongside VAF. Variants absent or observed in only one or two datasets were disproportionately enriched in difficult and extreme regions, a bias that became more pronounced at higher VAFs (**Fig. 2d–e**). By contrast, variants consistently observed across ≥3 datasets mirrored the expected genomic distribution (easy: 64–79%; difficult: 10–12%; extreme: 9–15%). These results demonstrate that alignment uncertainty in low-complexity regions remains a critical barrier to the reliable detection of ultra-low VAF variants.

### Generation of negative controls for accurate error estimation

We also established a negative control set to avoid misclassifying true variants as false positives (**Fig. 3a**) in the benchmark. Absent such a framework, undetected but potentially real mutations would confound benchmarking accuracy. For example, some loci carried alternative alleles in Illumina reads but fell below the thresholds required for variant calling; others were detected in Illumina but lacked PacBio support due to insufficient long-read depth; and still others represented subclonal variants in the pure cell lines that were missed during reference set construction. Failure to account for such cases would produce misleading performance estimates and unfairly penalize variant callers.

**Figure 3.**
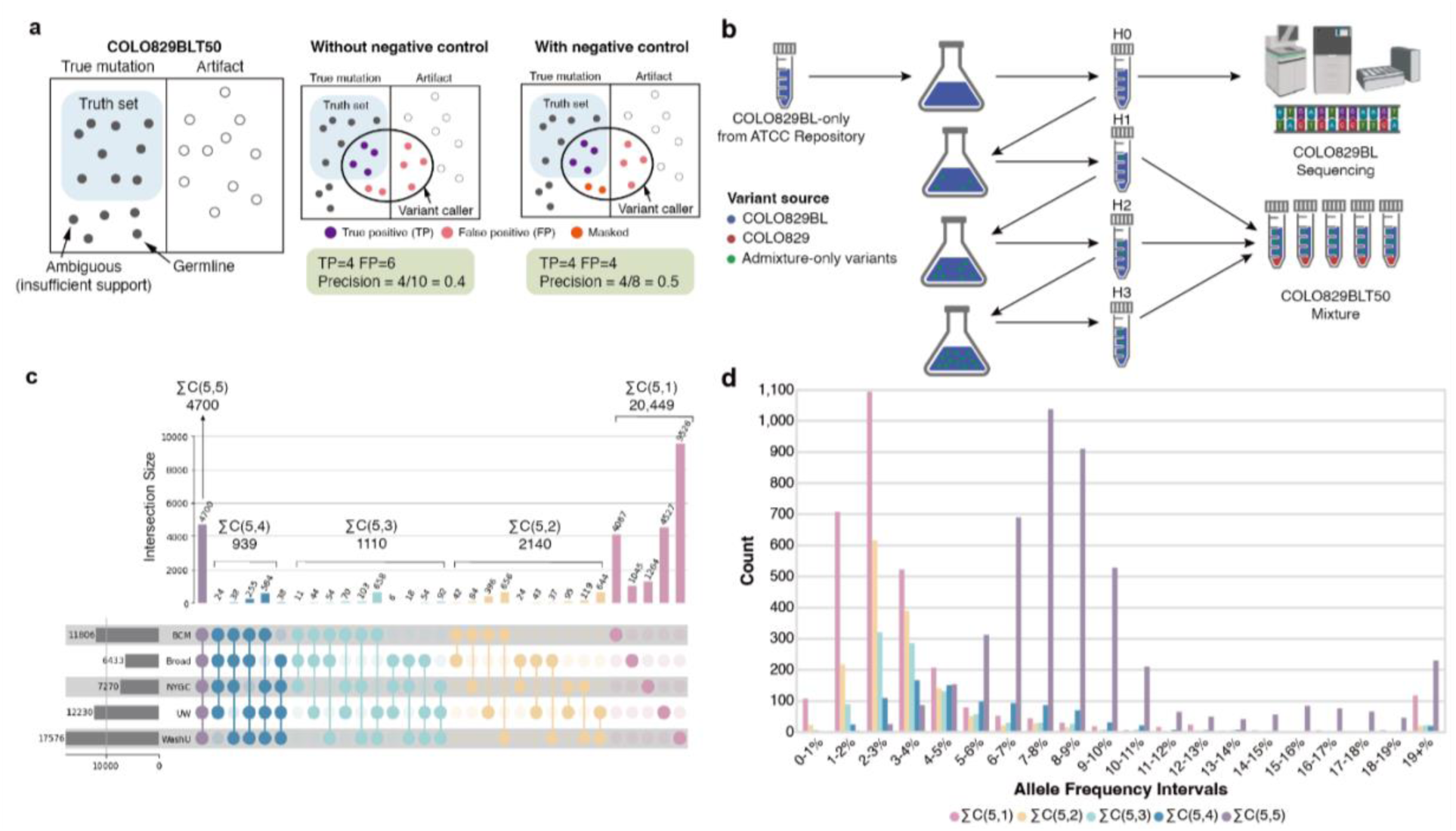
Importance of negative controls and evidence of cell culture-derived mutations. **a,** Conceptual schematic of the role of negative controls in variant benchmarking. Without a defined negative control set, true mutations that are not validated by PacBio data (possibly due to insufficient PacBio depth) may be incorrectly labeled as false positives, which, in turn, artificially reduces precision. Incorporating both positive (i.e., the reference set) and negative controls enables more accurate classification by separating artifacts from ambiguous but potentially real mutations. **b**, COLO829BL Harvest (H0) was sequenced prior to the subsequent cultures. The harvests, H1-H3, were used to create the COLO829BLT50 mixture, potentially allowing for emergence of somatic mutations in a normal lymphoblast cell line presumed to be stable. These mutations (visualized in green) were removed from the final reference set and masked from downstream analysis. **c,** The Upset plot shows all possible intersections of COLO829BLT50 admixture-specific mutations found across five COLO829BLT50 replicate datasets. **d**, The histogram shows distributions of allele frequencies for the summed intersection sets grouped in panel (c).

The negative control set comprised reference homozygous sites, defined as positions without alternate allele evidence in both COLO829 and COLO829BL (**Methods**).

Variants shared between the two cell lines, which are likely germline variants and unlikely to represent mosaic events in the COLO829BLT50, were also excluded. Positions that did not fall into either the reference set or the negative control set were annotated as *masked* and excluded from benchmarking owing to ambiguity in classification.

We categorized mSNV and mIndel sites into three groups:

1. **Reference set (positive control):** Mosaic variants detected using Illumina data and validated by PacBio data, present in COLO829 and absent in COLO829BL.
2. **Negative control:** Positions lacking alternate alleles in both cell lines or consistent with germline variation.
3. **Masked (ambiguous):** Sites not confidently assigned to either the reference or negative control sets, and thus excluded from evaluation.

### Identification of cell culture-derived mutations

Although cell lines are generally assumed to maintain stable genotypes across passages, immortalized cells can acquire *de novo* mutations over time, leading to subtle genomic divergence between harvests^13^. During assessment of putative false positives in the evaluation, we observed that many variants absent from the reference set exhibited biological features consistent with genuine mutations rather than technical noise, appearing at reference homozygous sites in both pure COLO829 and COLO829BL lines. In addition, mutational signature analysis of these variants in COLO829BLT revealed a strong ROS-associated signal (COSMIC signatures SBS18/17b). These signatures, already associated with artifactual mutations in cell cultures, were absent from in silico mixtures (**Figure S2**), indicating that some apparent false positives may instead reflect genuine culture-acquired mutations.

In preparing the COLO829BLT50 mixture, the normal cell line (COLO829BL) was harvested at four time points (H0–H3) with minimal passaging and was assumed to be genomically stable (**Fig. 3b**). The H0 harvest was used to define the COLO829BL genome for reference set construction, whereas H1 through H3 were used for generating the mixture datasets. This design left open the possibility that subclonal mutations arising during COLO829BL passaging–present in H1-H3 but absent from H0– would represent true mutations in the COLO829BLT50 mixture.

To capture potentially subclonal mutations and ensure a robust benchmark, we examined mutation calls detected in five biological COLO829BLT50 replicates but absent from the COLO829/COLO829BL-defined reference set (**Fig. 3c**). Of these, 4,699 calls were observed in all five replicates, with 939, 1,110, 2,140, and 20,449 detected in four, three, two, or one replicate, respectively. In examining the VAF distributions of variants found in all five COLO829BLT50 replicates, we found the largest number of variants in the 7-8% range (**Fig. 3d**, purple bars). Many of the variants found in a subset of the replicates were at or below 3% VAF (**Fig 3d**, pink bars), suggesting more subclonal mutations sampled and emphasizing the need for subsequent corroboration by orthogonal sequencing methods or experimental validation. We experimentally validated three putative culture-acquired mutations consistently found across all five replicates using droplet digital PCR (ddPCR), which confirmed their increasing allele frequencies during COLO829BL passages and their presence in the final COLO829BLT50 mixture (**Fig. S3**).

To comprehensively characterize putative COLO829BLT50 admixture-only mutation sites, we cross-referenced each site with PacBio data for COLO829BLT50 (387× coverage). Applying a minimum requirement of two supporting PacBio reads (**Fig. S4**), we identified 12,230 admixture-only mutations with long-read support (**Fig. S4b**, top bars). Because sequencing or systematic errors in the PacBio dataset could spuriously confirm false signals, we additionally compared these sites from unrelated control (∼200× coverage) and found that up to 5% were erroneously supported (**Fig. S4b** bottom bars, see **Methods**). Therefore, we refined the set to include only sites with ≥2 PacBio reads in COLO829BLT50 and no mutant allele support in the control and the two pure cell lines, yielding 5,532 high-confidence admixture-only mutations. These sites were incorporated into the COLO829/COLO829BL reference set, annotated as culture–derived sites. Together, these analyses underscore that robust benchmarking cannot rely on pure cell lines alone. Whereas most prior cell line–based benchmarks overlooked culture artifacts and ambiguous sites, our framework integrates curated positive and negative controls and explicitly validates uncertain loci. Such rigor is essential for ensuring that benchmarking truly reflects caller performance rather than hidden biases of the model system.

### The SMaHT low-VAF (<5%) Mosaic Mutation Detection Challenge

To assess the effectiveness of current methods for low-VAF mosaic SNV and Indel detection, we conducted a Network-wide benchmarking challenge. This effort focused on testing somatic variant detection pipelines applied to single-sample Illumina whole-genome sequencing data from the COLO829BLT50 mixture, which contained mosaic mutations with VAFs below 5%.

Five biological replicates of the mixture, sequenced at depths ranging from 180× to 500×, were used to represent real-world whole-genome data with biological and technical variability (**Table S1**). In addition, five *in silico* COLO829BLT50 datasets were generated by mixing BAM files of the parental cell lines at defined total depths from 100× to 500×, enabling a controlled evaluation of depth-dependent performance.

A total of seven institutions participated, contributing eleven variant calling approaches for both mSNVs and mIndels (**Methods**). These pipelines included tools developed specifically for mosaic variant detection or adapted from frameworks originally developed for somatic mutation detection in cancer (**Table 1 and Table S3**): MosaicForecast^14^, a machine learning model trained on read-level features, was included based on its previously reported high performance in exome-based benchmarks; Mutect2^15^, a widely used somatic variant caller, was run in tumor-only mode under three different configurations, varying post-calling filters based on VAF thresholds and depth cutoffs; Lancet^16^, a local-assembly-based method, was also tested for its ability to resolve complex or low-VAF mutations; and RUFUS^17^, a reference-agnostic sequence comparison tool, was evaluated for its alignment-free detection strategy. Although the challenge focused on single-sample variant calling, the limited use of the COLO829BL control sample was permitted for RUFUS. This allowance enabled evaluation of its sequence-based comparison algorithm, which bypasses alignment and may offer advantages in detecting mosaic variants in complex genomic regions.

**Table 1.**
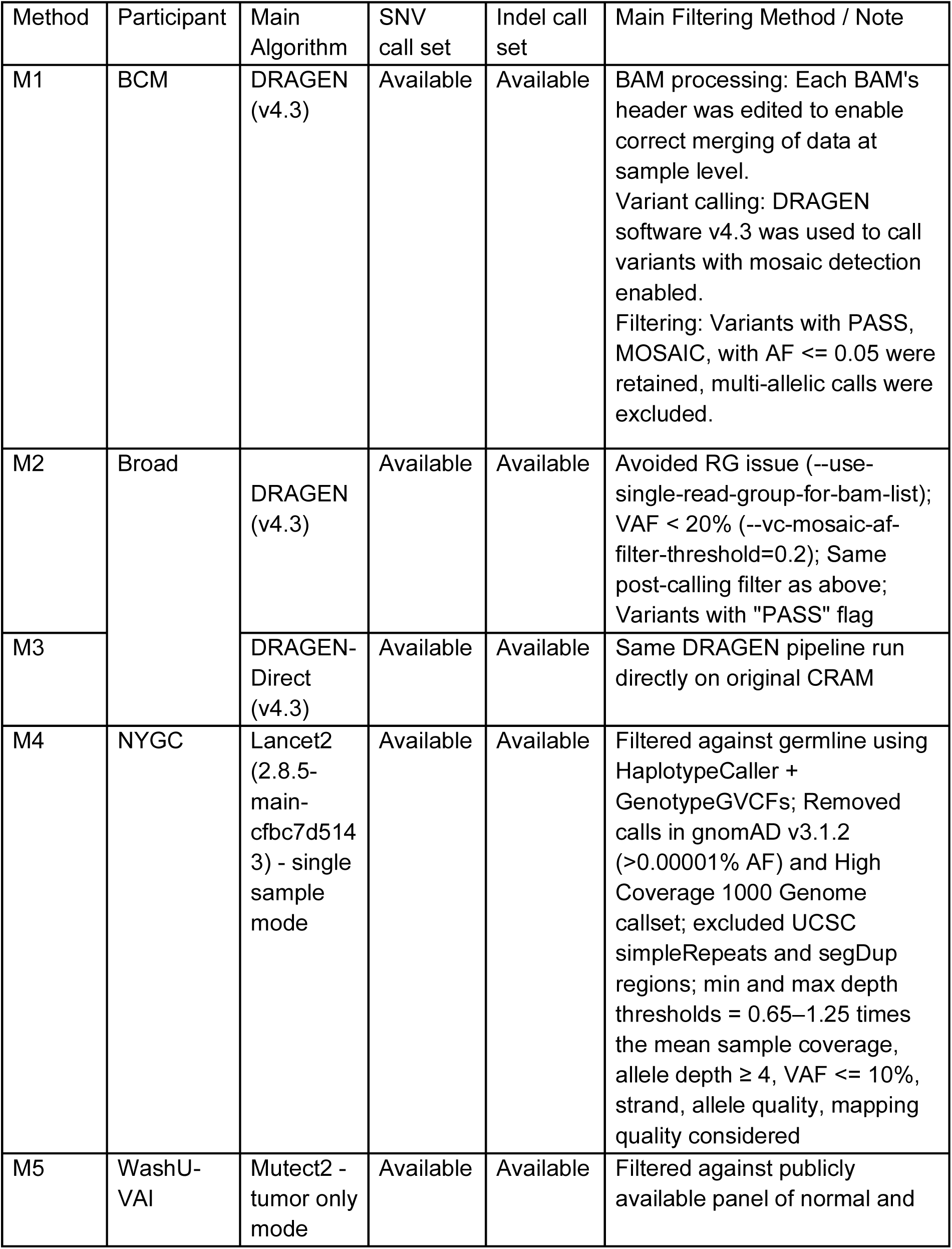
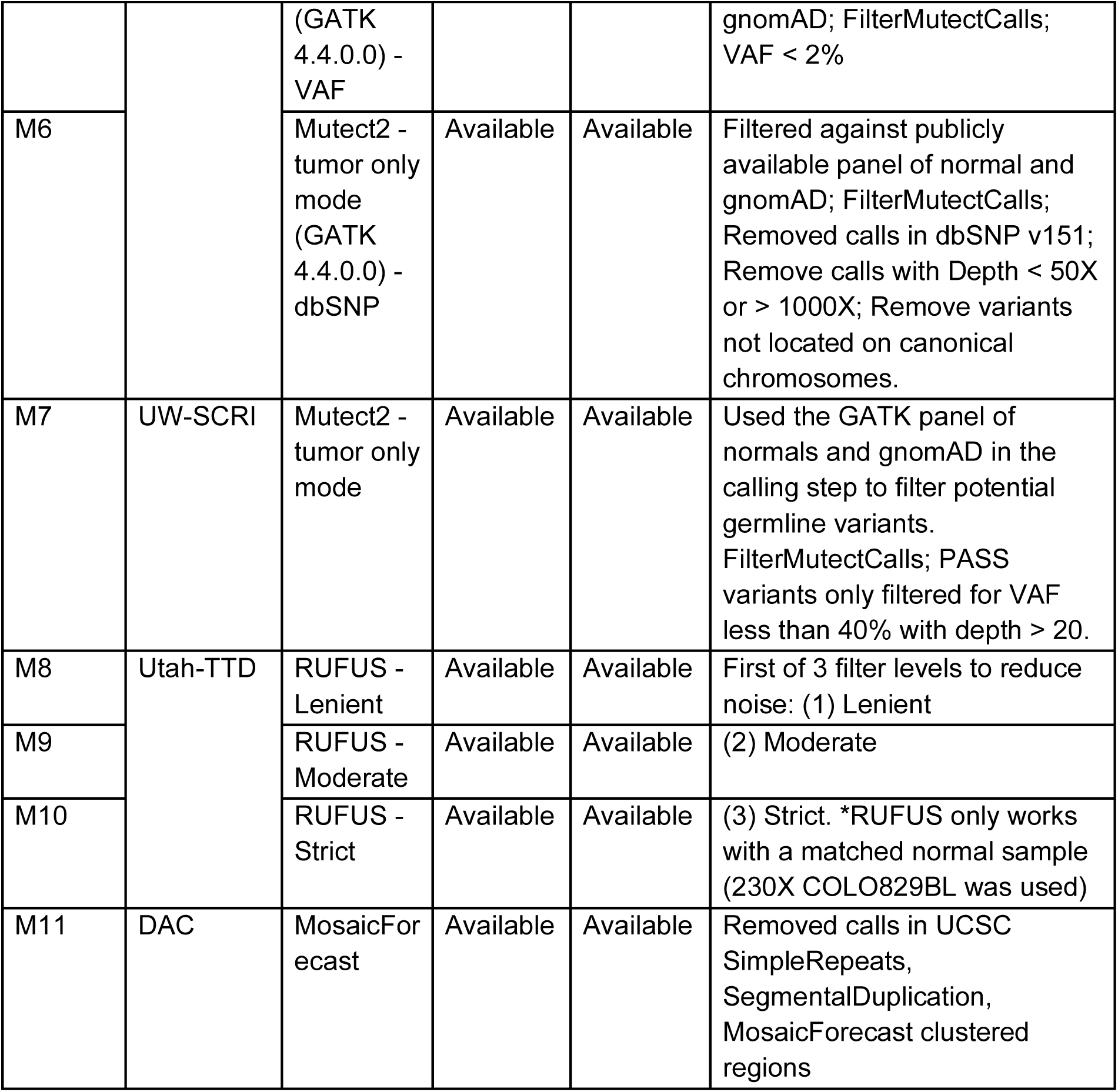
Eleven single sample-based mosaic mutation detection methods participated in the low-VAF Mosaic Mutation Detection Challenge.

### Performance evaluation of the detection challenge

Systematic evaluation across sequencing depths is particularly critical for low-VAF variants (<5%), where the number of supporting reads is inherently small (theoretically only 1 at 100× or 5 at 500×). In practice, such signals are further obscured by sequencing inconsistencies and low-quality features such as short read lengths, base-calling errors, and reduced base quality scores that exacerbate detection.

In the COLO829BLT50 mixtures, sensitivity of mSNVs was uniformly poor at 180×, with only a few pipelines reaching ∼50% in the 2–3% VAF bin (**Fig. 4a**). At 400×, nearly half of the methods exceeded 70% sensitivity, with MosaicForecast (M11), UW-SCRI Mutect2 (M7), and RUFUS with matched control (M8-M10) performing best. Pipelines enforcing hard VAF cutoffs, such as WashU-VAI Mutect2 (M5) and BCM DRAGEN (M1) with strict VAF filters (<0.02 and <0.05, respectively), showed marked limitations.

**Figure 4.**
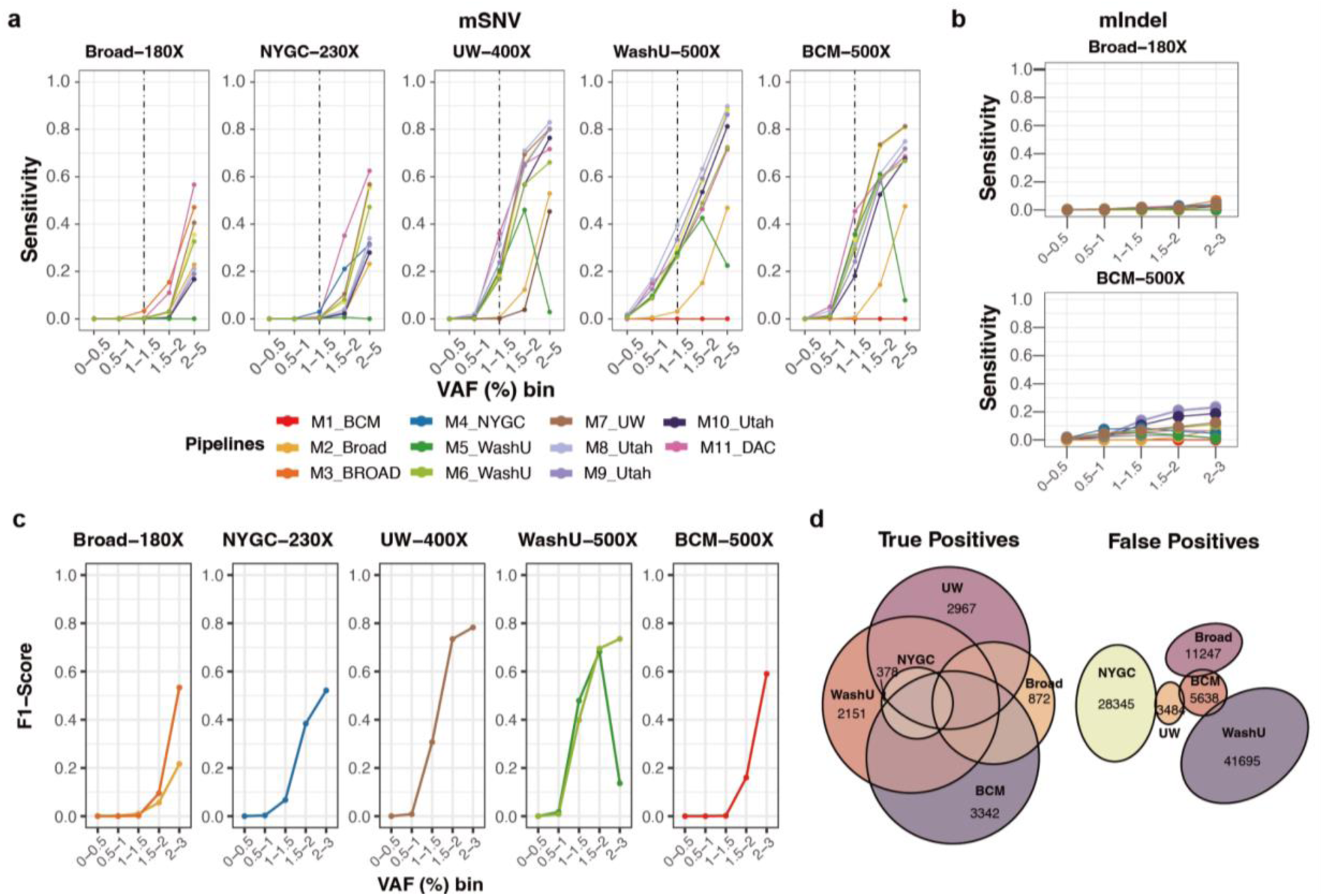
Performance benchmarking of low-VAF mosaic variant detection. **a,** Sensitivity across VAF bins for 11 detection pipelines applied to five biological COLO829BLT50 datasets (180×–500×). Methods are basically single-sample approaches, with RUFUS as the only matched-control approach. **b,** Sensitivity of mIndels across VAF bins for two representative depths (Broad 180× and BCM 500×). **c,** F1 scores stratified by VAF bin across five independently generated datasets, each analyzed using a distinct institutional pipeline, reflecting the combined effects of data quality and detection method on overall performance. **d,** Overlap of true positives (left) and false positives (right) detected from the five methods in panel c.

Because VAF is inherently depth-dependent and shaped by dataset-specific noise, such fixed thresholds often suppressed true positives. Precision at lower depth (180×) was poor in even 2-3% VAF, where most of the methods could recover high precision at VAF > 1% from 400× (**Figure S5**).

Among all evaluated tools, MosaicForecast provided the most balanced performance in terms of sensitivity and precision up to 230×. At 500×, probability-based methods such as Mutect2 gained greater statistical support for true alleles and outperformed other approaches.

To further assess the effect of sequencing depth, we applied the same pipelines to five *in silico* COLO829BLT50 mixtures generated by combining pure cell line BAMs at defined depths (100×–500×) (**Methods**). Similar patterns were observed: sensitivity for variants with VAF <1% remained poor and did not improve with higher depth. Gains in F1 score, reflecting the balance of sensitivity and precision, were evident up to 300× for VAF >1%. For example, the F1 score of MosaicForecast (M11) increased from 0.15 at 100× to 0.48, 0.68, and 0.75 at 200×, 300×, and 400×, respectively, but plateaued at 0.74 at 500× (**Figure S6** and **Table S4**). Overall, F1 scores improved steadily with increasing depth, although sensitivity remained the principal bottleneck, particularly for ultra-low VAF (<1%) variants across all pipelines.

For ultra-low-VAF mosaic Indels, sensitivity remained low even at 500×, reaching only ∼20% with RUFUS when applied with a matched control (**Fig. 4b** and **Figure S7-8**). Single-sample approaches achieved ∼10% sensitivity but with higher precision, reflecting a trade-off between recall and specificity. Given the biological importance of mIndels at low VAFs and the inherent mapping challenges of short reads spanning insertions or deletions, improved methods specifically designed to overcome this limitation are urgently needed.

We next evaluated a more practical scenario in which five independent groups (BCM, Broad, NYGC, UW-SCRI, and WashU-VAI) applied their own pipelines to independently generated COLO829BLT50 datasets. This design mimics real-world mosaic variant calling, enabling assessment of how variant profiles converge or diverge when the same biological material is analyzed across institutions for ultra-low-VAF mosaic mutations.

Performance largely tracked with sequencing depth (**Fig. 4c**), although some relatively lower-depth datasets-such as the 400× dataset from UW-SCRI-achieved F1 scores comparable to higher-depth datasets, underscoring the importance of both data quality and analysis strategy. Detected mutations were also highly distinct across groups, with thousands of true positives unique to individual pipelines (**Fig. 4d**). False positives varied substantially, with little overlap across methods. These results suggest that intersecting multiple callers may reduce false positives but at the expense of discarding many true variants, highlighting the need for more sophisticated algorithms for ultra-low-VAF mutation detection.

### Low-VAF mutation detection in different genomic regions

Lastly, we examined how genomic context influences the detection of low-VAF mutations by using high-confidence variant sets comprising 34,302, 5,216, and 4,487 SNVs and 591, 686, and 782 indels in easy, difficult, and extreme regions, respectively. To first assess whether VAF estimates differ between sequencing platforms, we compared Illumina and PacBio data across regions. VAFs for mSNVs were highly concordant in easy (*R* = 0.99, *p* < 10^-15^) and difficult (*R* = 0.98, *p* < 10^-15^) regions, but correlation was reduced in extreme regions (*R* = 0.92, *p* < 10^-15^) (**Fig. 5a**). Indels showed lower overall concordance, with correlations of 0.97, 0.84, and 0.71 in easy, difficult, and extreme regions, respectively. These results indicate that even in highly clonal COLO829 samples where mutations typically have VAFs near 50–100%, alignment ambiguity persists in short-read WGS. Such effects are expected to be exacerbated in mosaic contexts such as COLO829BLT50, where low-VAF mutations are intrinsically limited by lower signal-to-noise ratios, further complicating accurate detection.

**Figure 5.**
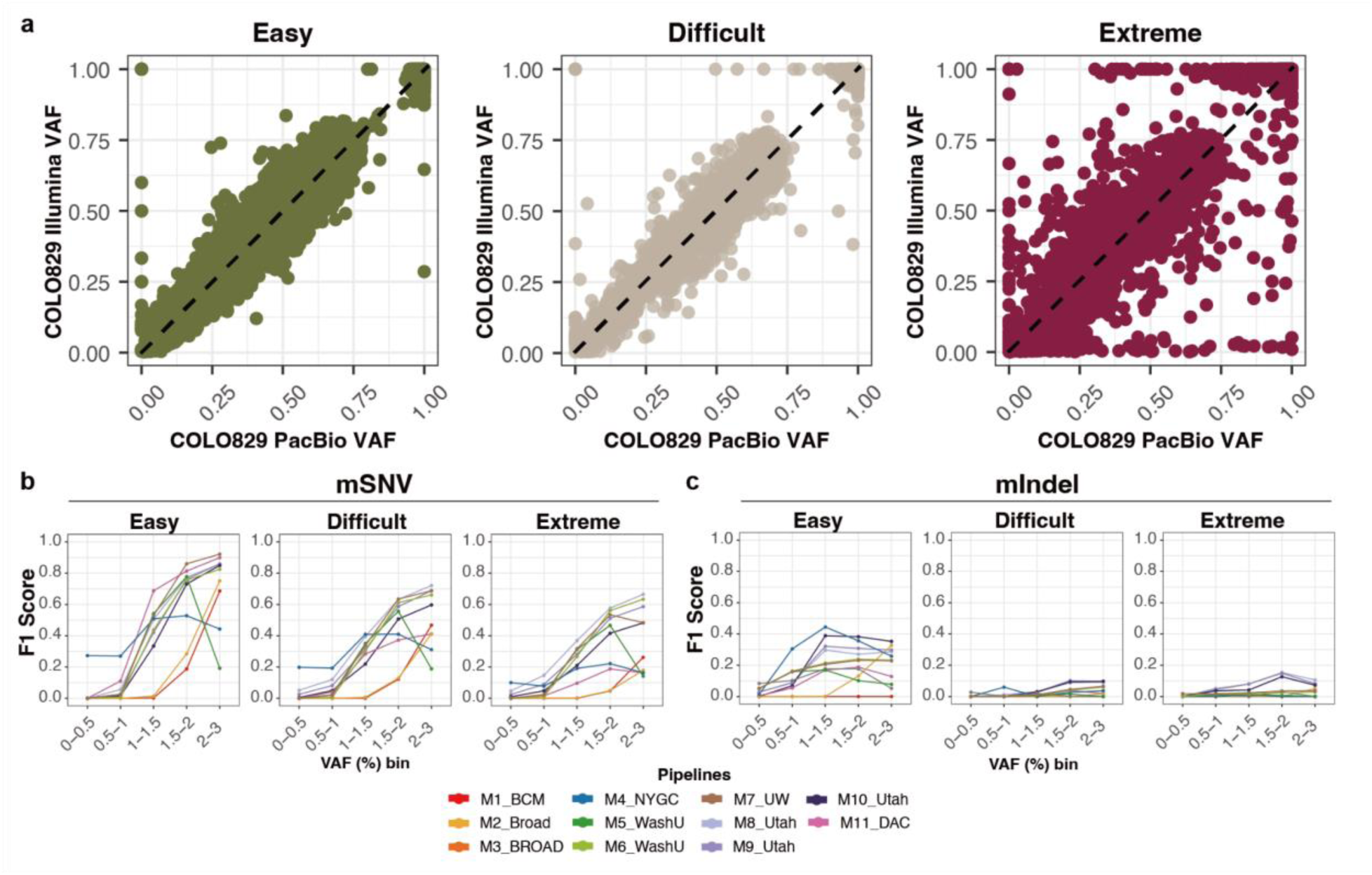
Impact of genomic region complexity on low-VAF mutation detection performance. **a**, VAF comparison between PacBio and Illumina for COLO829 across the three different genomic regions. Each point represents a validated mosaic SNV; color indicates region category (easy, difficult, extreme). **b-c**, F1 scores stratified by different genomic contexts (Easy, difficult, extreme). 500× BCM-generated COLO829BLT50 was used with ten detection pipelines. (b) mSNV and (c) mIndels.

We next evaluated detection performance across genomic contexts. For mSNVs in easy regions, 1% VAF mutations achieved ∼70% sensitivity and >90% precision, with F1 scores exceeding 0.8 for variants ≥2% VAF (**Fig. 5b** and **Figure S9**), consistent with prior exome-based benchmarks^7^. In contrast, sensitivity declined sharply in difficult and extreme regions: at 1% VAF, F1 scores dropped to ∼0.4, roughly half of those observed in easy regions. Methods generally clustered into two categories: those favoring precision (e.g., DRAGEN [M2-M3], MosaicForecast [M11]) and those favoring sensitivity (e.g., Mutect2-based pipelines [M5-M7]). Interestingly, single-sample methods (MosaicForecast [M11] and Mutect2 [M7]) outperformed matched-control approaches (M8-M10) in easy regions, suggesting that robust statistical or machine-learning models can distinguish true low-VAF mutations from artifacts without external controls.

In contrast, mIndel detection remained substantially more challenging. Alignment-free or alignment-modified methods yielded the best performance, with sensitivity in easy regions reaching only ∼0.4 at 1% VAF and rising to ∼0.6 at 3% when matched controls were used (M8-M10). Lancet (M4), a local assembly method, performed best at intermediate VAFs (0.5-1.5%), offering high precision with modest sensitivity. These findings underscore the urgent need for improved methods to address the alignment ambiguity of short reads at indel sites, as well as persistent sensitivity gaps for mSNVs in difficult and extreme regions. Overall, low-VAF mutation detection was constrained by artifacts that varied in impact across genomic contexts. By using long-read-supported reference sets, we were able to extend evaluation beyond easy regions and characterize the variant calling performance in difficult and extreme regions, where context-specific artifacts shaped distinct sensitivity and precision. These patterns provide valuable clues for developing more advanced methods for detecting ultra-low-VAF mosaic mutations across the whole genome.

#### Impact of sequencing depth and sampling variability based on HapMap cell line mixture

To further validate the consistency of analyses with COLO829BLT50 and to extend evaluation of detection limits beyond the 2% design of COLO829BLT50 up to ∼15% VAF, we employed a HapMap-based mixture model comprising ∼4.8 million reference SNVs. Six well-characterized HapMap cell lines (HG002, HG005, HG00438, HG02257, HG02486, and HG02622) were mixed to create target VAFs ranging from 0.25% to 16.5% (**Fig. 6a**). Although the six sample-mixture design does not permit haplotype phasing, it enables the evaluation of a larger number of variants across a broad range of VAFs (**Figure S10a**). The reference set for the HapMap mixture benchmarking experiment was created from curated germline SNV calls from Genome in a Bottle and the Human Pangenome Reference Consortium (**Methods and Table S5**). We generated a reference SNV set comprising ∼4.8M SNVs with about a half (2.1M) at VAF ≤1%, with multi-allelic sites removed and restricted to high-confident regions. HapMap mixture samples were sequenced up to ∼500× short-read and ∼100× long-read whole-genome coverage across four centers each (**Fig. 6b** and **Table S1**).

**Figure 6.**
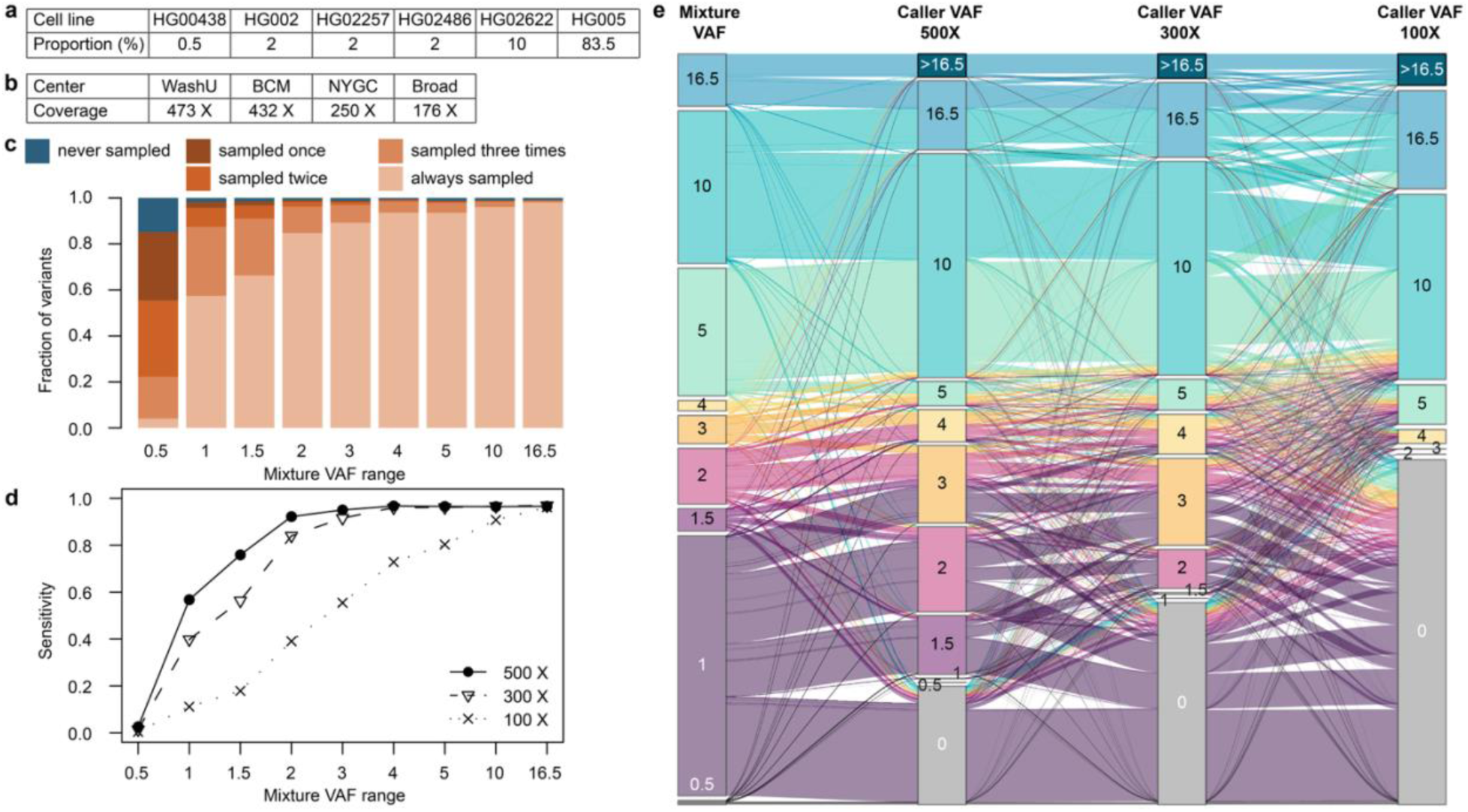
HapMap mixture analysis on sampling bias, sequencing depth, and VAF dynamics. **a**, Proportion (%) of cell lines in the HapMap mixture sample. **b**, Average sequencing depth of genome-wide coverage of Illumina short read data generated from technical replicates of the HapMap mixture at different centers. **c**, Sampling of somatic variants depends on their VAF. While >85% of variants with VAF ≤0.5 and >98% of variants with VAF in (0.5, 1] were sampled at least once in the 4 short read datasets, only 4% and 57% of variants were consistently sampled in all 4 datasets, respectively. With higher VAFs, variants are more reliably sampled in all datasets. **d,** Sensitivity of variant calling depends on sequencing depth, with an increasing number of low VAF variants called with deeper sequencing of 500× compared to 300× and 100× approaches. Subsampled 300× datasets showed comparable sensitivity to datasets aggregated to 300× from multiple 100× datasets subsampled from different datasets. **e,** Comparison between expected “mixture VAF” based on the proportion of variants in the HapMap mixture, and “caller VAFs” calculated by a variant caller, i.e., MuTect2, across different sequencing depths. Variants are colored by their mixture VAF range. At lower sequencing depths, the caller VAF deviates further from the mixture VAF, and false negative calls increase (indicated by gray bars labeled with 0).

For variants with expected VAFs > 1%, over 98.8% were observed at least once out of four short-read datasets of the HapMap mixture, and 92% could be consistently observed across all four replicates, reinforcing that even 200× coverage provides robust sensitivity (**Fig. 6c**). By contrast, only 4% of variants with up to 0.5% expected VAF were consistently sampled across all samples, but 85% were identified at least once, predominantly by the ∼500× samples.

Building on these sampling rates, we evaluated variant calling with Mutect2^15^ to assess how coverage and allele support translate into final mutation calls (**Figure S10c**).

Notably, sensitivity was improved substantially between 100× and 300× from 11% to 40% for 1% VAF variants and 39% to 84% for 2% VAF variants, but the gain is minimal beyond 300× (**Fig. 6d)**. This quantification of diminishing returns in variant detection sensitivity beyond 300× informed the SMaHT Network’s decision to adopt 300× as the target depth for all bulk sequencing. To evaluate the impact of merging sequencing datasets across centers, we combined three independent 100× subsampled datasets from different centers to achieve an aggregate 300× coverage dataset and compared variant calling performance with subsampled 300× datasets from individual datasets.

Sensitivity was nearly identical to that of single-center 300× datasets (**Figure S10c**), supporting the feasibility of merging data across centers to achieve higher coverage without loss of performance.

Because the mixture proportions were precisely defined, we were also able to compare VAF expectations with VAFs observed and reported by somatic variant callers at different sequencing depths (**Fig. 6e** and **Figure 10b**). In general, the VAF reported from Mutect2 (i.e., caller-VAF) was consistently higher than expected, reflecting sampling bias: variants that happen to receive more read support are more likely to be called. Overall, for 23% of variants the caller-VAF remained consistently within the expected mixture VAF range. However, most of these were high-VAF variants; only 4.7% of variants with expected VAF ≤5% were concordant, indicating that VAFs were often inflated, particularly at low frequencies. At 100× coverage, only 38 variants were called at the lowest reported VAF of 2%, with expected values between 0.5% and 4%, highlighting the limited precision of VAF determination for low-VAF somatic variants. This bias underscores the need for caution when interpreting VAFs from variant callers in applications where accurate quantification is critical.

## Discussion

Comprehensively detecting mosaic mutations remains one of the most difficult challenges in genome analysis, in part because it requires pushing the limit on distinguishing low-frequency mutational signals from the inherent background noise of high-throughput sequencing data. Although the cost of whole-genome sequencing has dropped significantly in recent years, simply increasing sequencing depth to improve the signal-to-noise ratio is not a cost-effective strategy. For example, even at 500× coverage, sensitivity for detecting mSNVs at 1–2% VAF remains 0.5, and it drops precipitously for variants at lower VAFs. This highlights a clear point of diminishing returns, where further increases in depth yield only marginal gains and fail to capture mosaic mutations beyond those arising in the earliest cell divisions.

We generated an exceptionally high-coverage whole-genome dataset to assess the impact of sequencing depth, variability in replicate experiments, and algorithmic performance on mosaic variant detection. Despite applying four Illumina-based methods for candidate identification, some variants were inevitably missed due to imperfect sensitivity of the current algorithms. To address this, we introduced a negative reference set and incorporated carefully characterized cell culture-derived mutations, enabling more robust evaluation of low-VAF mosaic mutations-features largely overlooked in previous cell line-based benchmarking designs.

Our results revealed that many Illumina-derived variants were not validated by PacBio reads, identifying important sources of false positives in Illumina-only pipelines. Some false positive calls were, in fact, rare germline variants in copy number-amplified regions-for example, a germline variant in a region with four copies can appear as mosaic (VAF ≈ 0.25) under the diploid assumption. Additional false positives arose, for instance, in segmental duplication or repetitive regions, where alignment ambiguities cause reads from multiple similar regions to collapse onto a single locus, making mutations in one copy appear as low-VAF mosaic events.

Notably, we generated high-coverage PacBio HiFi long-read WGS data, enabling confident orthogonal validation of candidate variants, including those in repetitive or difficult-to-align regions. This dual-platform approach significantly improved the quality of our curated reference set and revealed key limitations of benchmarks based solely on short-read data, demonstrating that artifacts impact sensitivity and precision differently across genomic contexts. Future methods will need to explicitly model and filter such artifacts, and the benchmarking resources provided here establish a foundation for developing and validating next-generation mosaic variant callers.

Although PacBio data were essential in building our reference set, obtaining high-depth PacBio HiFi data from tissues can be challenging in practice due to the requirement for high molecular weight DNA, which cannot be reliably extracted from many tissue types. However, our cross-platform validation has informed the development of more effective Illumina-only detection strategies, improving both accuracy and feasibility in diverse biological samples.

To overcome the limitation of high-depth whole-genome sequencing, complementary technologies are essential. Single-cell whole-genome sequencing, for instance, has been used to identify rare mutations, which can then inform the design of targeted panels for ultra-deep sequencing (e.g., >100,000×) across multiple tissues. Duplex sequencing, which enhances specificity by capturing both DNA strands of a molecule, offers another powerful strategy for detecting rare mosaic variants at the single-molecule level. The disadvantage of duplex sequencing, however, is that one cannot determine whether any two mutations have occurred in the same cell.

Another promising avenue involves leveraging multiple tissues from the same individual. In the SMaHT Network, up to 15 tissues per donor are sequenced when available. Early developmental mutations are expected to be shared across multiple tissues, while tissue-specific or germ layer-specific variants are likely confined to a subset. This tissue distribution can be exploited to improve variant filtering-for example, by using one tissue as a control for another in paired-calling frameworks to distinguish true tissue-specific variants from technical artifacts.

In summary, our study highlights the current limitations of using sequencing depth alone to detect low-VAF mosaic mutations using bulk short-read sequencing data and underscores the need for integrative approaches-combining ultra-deep sequencing, long-read validation, multi-tissue analysis, and advanced computational methods. The pure cell lines, cell line mixtures and curated reference sets developed here are publicly available and will serve as valuable resources for the community to benchmark and advance mosaic and low-VAF variant detection tools.

## Methods

### Sample preparation and data processing

#### WGS data acquisition, alignment, and QC

Illumina short read whole genome sequencing (WGS) data analyzed for COLO829 and COLO829BL can be obtained from the SMaHT Data Portal (https://data.smaht.org/). For the COLO829BL WGS data specifically, three BAM files, each derived from a sequencing library from the same COLO829BL analyte, were analyzed (file accession IDs: SMAFIARM7511, SMAFIJ9GV6FG, and SMAFIH6KN5FI). The sample preparation for all samples described in this study, sequencing, data processing, and alignment are described in detail in the SMaHT Network paper^18^. In brief, all WGS data were uniformly processed using standard pipelines for each sequencing platform: Illumina paired-end WGS data were processed using a pipeline based on the Genome Analysis Toolkit (GATK) Best Practices^19,20^. After raw sequence quality assessment, sequence artifacts are removed using fastp^21^. Reads were then aligned to the GRCh38 reference genome that excludes ALT contigs and decoy sequences using BWA-MEM^22^ (*v0.7.17*), followed by uniform post-processing steps for data quality assessment, including duplicate reads marked using Picard^23^, local realignment, and base alignment quality and base quality score recalibration (BQSR). PacBio HiFi WGS data were aligned to the same reference genome using pbmm2^24^ (*v1.13.0*), preserving methylation-related MM and ML tags in the final BAM files. All WGS data were evaluated for data quality and sample integrity checks (e.g., sample swap or duplicates, contamination checks) using standard tools such as FastQC^27^, Samtools^28^, and NanoPlot^29^, as well as in-house scripts implemented at the SMaHT Data Analysis Center.

### COLO829-BLT50 reference set generation

#### Illumina short-read-based candidate selection

To identify somatic SNVs and indels unique to COLO829, we applied four somatic variant callers, TNHaplotyper2^30^(*Sentieon Genomics 202112.06*), Strelka2^31^ (*v2.9.2*), VarNet^32^ (*v1.1.0*), and RUFUS^33^ (*v0.1*), using COLO829BL as the matched control. Each caller was executed with its recommended settings with 30Mb intervals and accompanied by internal variant filtering. For Strelka2, small indels were obtained using Manta^34^ (*v1.6.0*). To ensure consistent representation of variants across tools, all outputs were normalized with bcftools^35^ (*v1.14*; *norm -a -m - -f*). Only *PASS* calls were used and Indels were restricted to those <50 bp in length.

#### Collecting mutation-supporting reads from PacBio long-read

Variant support from PacBio HiFi data was assessed using bcftools mpileup^35^ *(v1.17)*, applying filters of maximum depth 3,000, minimum base quality 20, minimum mapping quality 1, and excluding supplementary, secondary, duplicate, and QC-failed reads. Resulting pileups were normalized with bcftools *norm* (allele splitting and left alignment) to generate per-chromosome PacBio pileup VCFs for both COLO829 and COLO829BL. For Indels, we developed a custom Python script (*Python v3.9.7, pysam*^36^ *v0.21.0*) to annotate indel-supporting reads from PacBio or Illumina BAMs at variant positions specified in VCF files. For each Indel record, the script fetches aligned reads from the BAM, filters by mapping quality (≥1 by default), and classifies reads as supporting the reference allele, the targeted alternative allele, or other alternative alleles based on sequence alignment at the breakpoint. The script calculates allele frequency (AF), read depth (Depth), supporting reference reads (DR), supporting alternative reads (DTV), and allele counts of non-target alternative sequences (*OtherAlt*), and appends these fields to the *INFO* column of the output VCF.

#### Validation with PacBio long-reads

To validate short-read variant calls, we developed Python pipelines (Python *v3.9.7*, pysam^36^ *v0.21.0,* SciPy^37^ *v1.11.3*) that cross-referenced Illumina-derived SNVs and indels with long-read PacBio pileup support. For SNVs, alternate allele counts (AD) from PacBio tumor and blood BAMs were parsed per variant, and calls were retained if supported by ≥2 tumor reads, absent from the matched blood sample, and not superseded by a competing allele at multi-allelic loci. Multi-allelic variants were resolved by selecting the major alternate allele based on tumor read counts. For Indels, additional annotation of alternate and non-target alleles (OtherAlt) was performed to quantify allele balance. Candidate Indels were tested for allelic imbalance using a binomial test (p > 0.05) and retained if supported in PacBio and Illumina tumor data (VAF > 0.2) but absent from Illumina blood (VAF < 0.05). Insertions and deletions ≥51 bp were excluded.

### Negative control construction for COLO829BLT50

#### 1) Nonvariant sites

To define the negative control set, we first identified homozygous reference sites shared by COLO829 and COLO829BL Illumina WGS. To define potential non-variant positions shared by COLO829 and COLO829BL, Santieon Haplotyper^30^ in the GVCF mode was applied independently on the two samples and putative homozygous positions shared by both were intersected for further processing. Next, we applied *bcftools mpileup* on the alignment of COLO829 and COLO829BL PacBio WGS in the putative homozygous candidates with the minimum mapping quality for an alignment (*q* parameter) and minimum base quality for a base to be considered (*Q* parameter) set to 1. Positions where no alternate allele found in both COLO829 and COLO829BL were collected as homozygous reference sites and included in the negative control set. The rest of the positions were then subjected to the next iteration of the *mpileup* analysis but with increased *q* and *Q* parameters (both set to 30). If alternate allele was observed with the higher quality threshold, those were removed from the negative control. Then, we also checked Illumina WGS data for to confirm remaining ambiguity. We again utilized *bcftools mpileup* on the short-reads with adjusted *q* and *Q* parameters (set to 1) and juxtaposed with the results of PacBio WGS pileups. Variants supported by both platforms were excluded, whereas those lacking Illumina support were retained as high-confidence negatives.

#### 2) Germline variants

We collected germline variants in COLO829BLwith Santieon Haplotyper^30^. Filtration involved several steps: First, GATK (*v4.1.9.0*) hard filters were applied based on GATK best practice recommendations (*QD* < 2.0, *FS* > 60, *DP* < 10, *ReadPosRankSum* < -8.0, and *MappingQualityRankSum* < -2.5 or > 2.5), retaining only PASS variants. Second, to remove germline-like systematic errors specific to PacBio data, particularly in homopolymer and tandem repeat regions, a binomial test [Binom(N, k, p) = Binom(depth, alternative allele count, 0.5)] with a p-value threshold of 0.05 was applied to distinguish heterozygous indels from subclonal Indels and artifacts.

### Characterization of COLO829BLT50 admixture cell-line artifacts

#### 1) Generating Admixture-Only Variant Sets

Artifacts of the COLO829BLT50 mixture were first suspected after comparing the RUFUS-reported allele frequencies from variants generated from the *in-silico* COLO829BLT50 data sets vs the physical admixture data sets. Within only the COLO829BLT50 admixture data sets, we observed variants reporting allele frequencies in the 5-10% range, i.e. allele frequency values seemingly inconsistent with the 1:49 tumor/normal blood cell line mixture. Manual inspection of several sites revealed the absence of the called mutant allele from the tumor sequencing dataset. To further investigate these variants, we examined call sets generated by both RUFUS and Mutect2 (performed in COLO829BLT50/BL paired calling mode) within each GCC-specific replicate COLO829BLT50 datasets. RUFUS and Mutect2 call sets within each replicate, were normalized and merged, creating a distinct call set within each of the five replicates. Any variants within these call sets that existed within the COLO828/COLO829BL reference set were removed, leaving only COLO829BLT50 admixture-specific variants. We then created 5-way intersections of these replicate call sets, as shown in the upset plot in Fig. 3c. We identified sites present in all five replicate datasets, i.e. C(5,5); then those present in 4, 3, 2, or only 1 of the 5 replicate datasets, i.e. C(5,4), C(5,3), C(5,2), and C(5,1), with the hypothesis that sites called within fewer replicate datasets represented lower confidence for the existence of the variant in the COLO829BLT50 mix.

#### 2) Characterization of COLO829BLT50 admixture-only variants in PacBio long-read datasets

To characterize each site, we performed a pileup in a 387× deep long-read dataset generated with the PacBio Revio sequencing instrument, and tabulated the number of PacBio reads supporting the mutant allele at the same site. We identified sites for which at least two supporting long reads were present. To avoid spurious confirmation caused by genomic site-specific sequencing errors, we repeated this procedure in a control sequencing dataset from a human lung tissue homogenate sample from an unrelated individual (ST001-1A), which was prepared using identical DNA extraction, library construction, and sequencing protocols to the COLO829BLT50 mix. We identified all sites at which the mutant allele called in the COLO829BLT50 dataset was supported by at least two PacBio long reads in the negative control dataset.

#### 3) Construction of the COLO829BLT50 admixture-only variant reference set

We have finally identified all COLO829BLT50 admixture-only candidate variants with a minimum of two supporting long reads in the COLO829BLT50 dataset, excluding sites with spurious “confirmation” in a control dataset. We have carefully identified the provenance of each reference-set variant whether it originated in the COLO829/COLO829BL-generated reference set or in the COLO829BLT50 admixture-only reference set. The identified sites were examined using both PacBio and Illumina alignments of COLO829 and COLO829BL. The datasets were queried using *bcftools mpileup* with filters of minimum mapping quality of 1 (*-min-MQ* 1) and minimum base quality of 20 (*-min-BQ* 20). Variants present in COLO829BL but absent in COLO829, with at least 1 read of support in either sequencing dataset, were categorized as COLO829BL-only mutations. Mutations with no evidence in either COLO829 or COLO829BL were intersected with the negative control regions to retain only high confidence variants forming the COLO829BLT50 admixture-only mutation reference set.

#### Mutational signature analysis

Mutational signature analysis was performed on false positive (FP) mutation calls annotated as “nonvariant”. A mutation count matrix was constructed by stratifying these FPs by sample, VAF bin, and variant calling pipeline. De novo mutational signatures were extracted from this mutation count matrix with MuSiCal^38^. Two of the discovered signatures – showing similarities to COSMIC SBS18 and SBS17b – were likely associated with ROS damages in cell cultures and shown in **Figure S2**.

### In silico COLO829BLT50 generation

The COLO829 and COLO829BL Illumina data used for reference set generation were downsampled with picard DownSampleSam (STRATEGY=HighAccuracy) with 2% and 98% respectively to the target depths: 100×, 200×, 300×, 400×, and 500×. The depths of the downsampled BAM files were measured with picard CollectWgsMetrics to confirm the accuracy of downsampling. Then, those BAM files were merged with samtools merge. For downsampling and merging, 30 Mbp intervals were used.

### Mosaic Mutation Detection Challenge pipelines

#### BCM

The Sentieon-generated GRCh38 BAMs for COLO829BLT50 sample-mix, were merged from each of the five sequencing centers. Variant calling was performed using Illumina’s Dynamic Read Analysis for GENomics (DRAGEN)^39^ software *v4.3*. Mosaic detection was enabled (*--vc-enable-mosaic-detection true*) with defaults used for other variant calling options. Variants on autosomes and allosomes tagged as PASS, MOSAIC, with *AF* <= 0.05 were retained using bcftools^35^ *v1.19;* any mult-allelic variant calls were excluded. *In-silico* datasets were not processed at BCM-HGSC.

#### Broad

Variant calling was performed using Illumina’s Dynamic Read Analysis for GENomics (DRAGEN)^36^ software *v4.3* for all cell-admixture and *in-silico* COLO829BLT50 samples. To ensure all reads were assigned to a single read group in BAM files of each sample, the *--use-single-read-group-for-bam-list* parameter was applied. Variants with VAF lower than 20% and tagged as PASS were retained. The same pipeline was applied on a single COLO829BLT50 sample sequenced at the Broad institute but without the pre-preprocessing of the read group information in BAM files.

#### NYGC

Mosaic SNVs and Indels were detected using Lancet2 (*2.8.5-main-cfbc7d5143*) in the single sample mode. The candidates were subjected to a series of following filters – Germline variants, identified with HaplotypeCaller and GenotypeGVCFs, were filtered out. Variants present in either gnomAD *v3.1.2* (with allele frequency greater than 1*10-5%) or in the high coverage NYGC generated 1000 genome call sets were removed. Variants in simple repeat (UCSC) and segmental duplication regions were excluded. Candidate sites were retained if they had a total depth between 0.65× and 1.25× of the mean sample coverage, with at least four supporting reads, and a variant allele fraction (VAF) ≤ 10%. Additional filters accounting for strand bias, base quality, and mapping quality were applied to retain only high-confidence mosaic SNV and Indel calls.

#### WashU-VAI

Somatic SNPs and indels were identified from COLO829BLT50 BAM files using GATK Mutect2 (*v4.4.0.0*) in tumor-only mode, with the GRCh38 reference genome, incorporating the gnomAD germline resource, and a 1000 Genomes panel of normal. To improve computational efficiency, the genome was divided into 30 intervals with *SplitIntervals*, and Mutect2 was run in parallel. Interval-level variant calls and statistics were merged using *MergeVcfs* and *MergeMutectStats*, and high-confidence variants were obtained with *FilterMutectCalls*. Two complementary post-calling strategies were applied. In the first, variants with allele frequency <0.02 were retained using bcftools, consistent with the expected 1:49 tumor–normal mixture (M5 in **Table 1**). In the second, variants were annotated against dbSNP151 using ANNOVAR^40^, and only novel SNPs and indels were retained (M6 in **Table 1**). This set was further filtered to exclude calls with sequencing depth <50× or >1000×, allele frequency >20%, and variants outside autosomes, chrX, chrY, and chrM. The resulting VCFs contained the final set of high-confidence somatic variants used for downstream analyses.

#### UW-SCRI

Bam files from the sequenced COLO829BLT50 resource from each GCC were analyzed with GATK Mutect2 in tumor only mode using a panel of normals and the gnomAD resource both downloaded from the Broad Institute to filter likely germline variants. Only *FILTER=PASS* variants were included using standard Mutect2 filters. The final call sets were filtered to require a depth >20 and VAF less than 40%.

#### DAC

MosaicForecast^14^ was run within the official MosaicForecast Docker container (https://hub.docker.com/r/yanmei/mosaicforecast). We first ran Sentieon TNHaplotyper2^30^ *v202308.03* (GATK Mutect2 equivalent) followed by Sentieon TNFilter *v202308.03*. To split multi-nucleotide and mutli-allelic variants into separate records, output VCFs were processed with bcftools^35^ (*v1.14*; *norm -a -m - -f*). Segmental duplication and variant clustered regions were removed from the VCFs before running MosaicForecast. The MosaicForecast *ReadLevel_Features_extraction.py* script was applied with the following parameters: 1) Umap mapability scores for k=24 generated based on the repository’s provided script, (2) candidate variants pre-formatted from VCF to BED, (3) the corresponding BAM file and (4) the reference genome. *Refine* models were selected based on the sequencing coverage of the input BAMs.

#### Utah-TTD

COLO829BLT50 bam files from a single GCC were initially merged for a single subject file input. RUFUS^33^ was run in a parallelized 1mb windowed mode, using COLO829BL as the paired control. Standard input parameters of a minimum k-mer threshold of 5 and k-mer length of 25bp were used, along with the ReportLowFrequency flag to allow identification of somatic mutations. After all 1mb region runs were completed, the mb variant call sets were trimmed to exclude any reported variants outside of the specified region and then concatenated into a single genome-wide VCF. A standardized RUFUS post-process script was applied (included in the tool Singularity container) and indels and SVs were removed to result in a final SNV-only, genome wide call set. RUFUS final VCF calls were evaluated using a variant quality score (VQS) classifier to remove potential false positive calls. The VQS model was a logistic regression trained on features from validated SNVs and indels from the SEQC2 HCC1395 tumor/normal benchmark dataset^41^. Three filtering stringencies were applied to the RUFUS call set. For the *lenient* threshold, no VQS filtering was applied. The *moderate* threshold corresponded to a VQS cutoff that achieved ∼90% sensitivity to true mutations for both SNVs and indels in the training data, while the *strict* threshold corresponded to cutoffs yielding ∼80% sensitivity for SNVs and ∼60% sensitivity for indels. These thresholds were designed to progressively reduce false positive calls while maintaining defined levels of sensitivity in mutation detection.

### Evaluation of Mosaic Mutation Detection Challenge pipelines

To harmonize variant representations, we used bcftools^35^ (*v1.14*) to split multi-nucleotide variants (MNVs) and multi-allelic sites into standardized single-allele records. For benchmarking, variant allele fractions (VAFs) were extracted directly from COLO829BLT50 BAM files using bcftools *mpileup* (base quality and mapping quality ≥ 1) for all mSNVs and mIndels in the reference set, including cell culture-derived mutations. If a reported mutations from pipelines were confirmed in the reference set, they were marked as true positives. If a mutation was found in negative controls, it was marked as false positive. If not found in both reference set and negative control, it was masked and not used for the evaluation due to ambiguity in the classification. For variants outside the reference set, VAFs were taken from the values reported by the respective callers. Sensitivity (truth positives out of all reference set), precision (truth positives divided by the sum or true and false positives), and F1 scores (balanced mean of sensitivity and precision) were computed across six VAF bins (0–0.5%, 0.5– 1%, 1–1.5%, 1.5–2%, 2–3%), chosen to distribute reference set variants more evenly across bins and enable fairer comparisons among methods.

### Evaluation across different genomic contexts

Easy regions were defined by the 1000 Genomes strict mask (high-confidence, short-read– callable regions). Difficult regions corresponded to sites within the PanMask^42^ strict (pm151b) set but outside the 1KG strict mask, representing moderately mappable regions. PanMask, a recent pan-genome-based accessibility mask derived from hundreds of high-quality assemblies, was used to provide a more generalizable and less reference-biased framework for benchmarking. Extreme regions encompassed loci outside PanMask, typically highly repetitive or structurally complex.

### HapMap mixture experiment

#### Experimental design

The SMaHT consortium designed several benchmarking experiments^18^ to evaluate somatic variant detection. One of these benchmarking experiments is a mixture of six clonal, well-characterized HapMap cell lines at different proportions, creating artificial somatic variants spanning VAF ranges below 1%, 1–2%, 5–10%, and above 10%. This mixture was intended to establish lower limits of detectability for sequencing technologies and somatic variant callers. Specifically, the mixture consisted of HG00438 at 0.5%, HG002, HG02257, and HG02486 at 2% each, HG02622 at 10%, and HG005 as background at 83.5%. Together, the heterozygous and homozygous germline variants, both private and recurrent across these cell lines (**Table S5)**, resulted in artificial somatic variants spanning VAFs from 0.25% to 16.5% (**Figure S10b**).

#### HapMap reference variant set construction

To generate a reference set of expected variants and their anticipated VAFs in the HapMap mixture, we first obtained germline variant call sets for the individual cell lines. Curated germline variants and sample-specific confident regions for HG002 and HG005 were obtained from the Genome in a Bottle (GIAB) benchmarking effort^43^, while those for HG00438, HG02257, HG02486, and HG02622 were obtained from the Human Pangenome Reference Consortium^44^. For all six cell lines, only SNVs within the intersection of the sample-specific confident regions were retained, covering 2.43 Gb of the genome. Indels, multiallelic variants, and SNVs with ambiguous reference bases (“N”) were excluded. Expected mixture VAFs were calculated based on the heterozygous or homozygous status of variants and the mixing proportions of the cell lines they occur in. In total, the HapMap mixture reference set contained 8,031,751 SNVs, with 4,888,479 within the artificial somatic VAF range (0.25–16.5%). Most of the somatic variants were private heterozygous variants from the 2% contributing cell lines, yielding >1,774,000 variants expected at 1% VAF.

#### Data generation

Bulk WGS datasets of the HapMap mixture were generated by multiple GCCs^18^. Illumina short-read WGS data were generated at 473× (WashU-VAI GCC), 432× (BCM GCC), 250× (NYGC GCC), and 167× (Broad GCC). PacBio long-read WGS data were generated at ∼100× (BCM, Broad, WashU-VAI GCCs) and 65× (UW-SCRI). See **Table S1**. All sequencing datasets were processed, aligned, and quality-controlled at the Data Analysis Center (DAC) as described in detail the SMaHT Benchmarking Publication^18^ and are available at the SMaHT Data Portal (https://data.smaht.org/).

#### Subsampling and merging of datasets

To assess the impact of sequencing depth on variant sampling and sensitivity of variant calling at low VAFs, short-read datasets were subsampled using *samtools view –subsample* ^35^(samtools *v1.17*) to 300× (WashU-VAI and BCM) and 100× (WashU-VAI, BCM, and NYGC). To evaluate whether aggregating lower-coverage datasets across GCCs could approximate higher-coverage sequencing by one GCC, all 100× subsampled datasets from different GCCs were combined to form a 300× aggregate dataset.

#### Variant allele sampling

To investigate the presence of expected variants in the HapMap datasets, we examined allelic support at all HapMap reference positions in both Illumina and PacBio datasets. Pileups were generated using *bcftools mpileup*^35^ (*v1.17*) with minimum base quality and mapping quality thresholds set to 30, ensuring only high-confidence bases in reliably aligned reads were considered.

#### Somatic variant calling in HapMap mixture

Many somatic variant callers were developed in a cancer research framework and are designed for tumor-normal comparisons. We applied callers with tumor-only functionality, given that the HapMap mixture mimics somatic variants without matched normals. Although the experimental design recapitulates low-VAF somatic variants, it differs from healthy human tissue samples in two aspects: the exceptionally large number of low-VAF variants (>4.8 million) and the presence of 12 haplotypes from six unrelated donors. To account for these design-specific aspects, we retained not only PASS variants but also those flagged for haplotype clustering (multiple nearby variants on the same haplotype) or variants annotated in germline datasets. Importantly, variants flagged with technical or quality-related filters (e.g., “base_qual,” “LowQual”) were excluded. Multiallelic variants were decomposed with *bcftools norm -m-any*, and indels or variants outside the confident regions were removed. Variants originating from HG005, the background cell-line, were considered germline and removed. False positives recurring across multiple samples were also excluded.

#### Illumina variant calling

Somatic variants in the Illumina datasets were called using *TNhaplotyper2* (Sentieon Genomics *202112.06*)^30^, the Sentieon implementation of GATK Mutect2, in tumor-only mode. Variants flagged as “clustered_events” (indicating multiple events on the same haplotype) or “haplotype” (variants co-occurring with filtered variants on the same haplotype) were retained alongside PASS calls.

## Data availability

The whole genome sequencing data generated by the SMaHT network (COLO829, COLO829BL, COLO829BLT50, and HapMap mixture) can be found at https://data.smaht.org/. The reference sets for COLO829BLT50 and HapMap mixture can be found at Github (https://github.com/parklab/SMaHT_SNV_COLO829BLT50_HAPMAP) along with negative controls and cell-culture derived mutations for COLO829BLT50. The reference genome GRCh38 (GenBank accession: GCA_000001405.15) is available at https://ftp.ncbi.nlm.nih.gov/genomes/all/GCA/000/001/405/GCA_000001405.15_GRCh38/seqs_for_alignment_pipelines.ucsc_ids/. Different genomic categorizations of easy, difficult, extreme regions can be found in the Github.

## Code availability

All pipelines and in-house scripts used for analysis can be found at https://github.com/parklab/SMaHT_SNV_COLO829BLT50_HAPMAP

## Acknowledgments

This research is supported by the NIH Common Fund, through the Office of Strategic Coordination/Office of the NIH Director under awards 1UM1DA058230, UM1DA058220, 1UG3NS132134, 1UM1DA058229, UM1DA058219, 1U24NS132103, UM1DA058235, UM1DA058236, UM1DA058237, and UM1DA058219.

## Author Contributions

P.J.P., Y-J.J.H., J.T.B., G.T.M., R.G., S.G., R.S.F., and K.A. supervised the project. Y-J.J.H. and D.M. generated the reference set and conducted all analysis for the COLO829BLT50 benchmark. J.M. performed analysis on HapMap mixtures with reference set generation. S.J.G. carried out cell culture-derived mutation analysis. N.L.P., T.C., S.D., S.G., D.K., D.L., B.M., R.M., L.X., Z.Y., and K.W., participated in the ultra-low VAF Mosaic Mutation Calling Challenge. L.A., N.Y.T.A., C.C., H.D., C.M.G., D.M.J., K.L., A.N., J.O., M.M.P, A.M.R., A.S., L.M.S., and T.W. contributed to biological material and data generation. H.J. performed signature analysis on COLO829BLT50. Y.Z. developed a pileup script for Indel analysis for short- and long-read data. C.T.K. contributed to regional stratification with Y-J.J.H.. M.B., W.C.F., and A.V. contributed to data processing at the SMaHT Data Analysis Center under the supervision of H-J.E.C. and P.J.P. Y-J.J.H. and P.J.P. drafted the manuscript with contributions from D.M., J.M., S.J.G. and G.T.M.

## Competing interests

Peter Park a member of the Scientific Advisory Board for Bioskryb Genomics. James Bennett is a consultant for Mosaica Medicines. Andrew Stergachis is a co-inventor on a patent relating to the Fiber-seq and DAF-seq methods.

Notes and Supplementary Figures

## Supplementary Notes

### Quality controls for sequencing data

For each BAM file, standard sequencing quality metrics, such as mapping rates, estimated insert sizes (**Figure S1**), average whole genome coverage, and other alignment-based statistics, were analyzed using Samtools^1^ (*v1.17*), Picard^2^ (*v3.0.0*), mosdepth^3^ (*v0.3.9*), and a custom in-house Python script. For PacBio and ONT data, reads were evaluated to assess the read-length distribution, N50, and other QC metrics specific to long-read data. For Illumina data generated on the NovaSeq platforms, raw reads were evaluated for base quality and base composition, and reads containing poly-G artifacts were removed using FastQC^4^ (*v.12.0*). The full description of the data quality control (QC) and assessment at the SMaHT Data Analysis Center is in the SMaHT Benchmark Flagship paper ^5^.

### Supplementary figures

**Figure S1.**
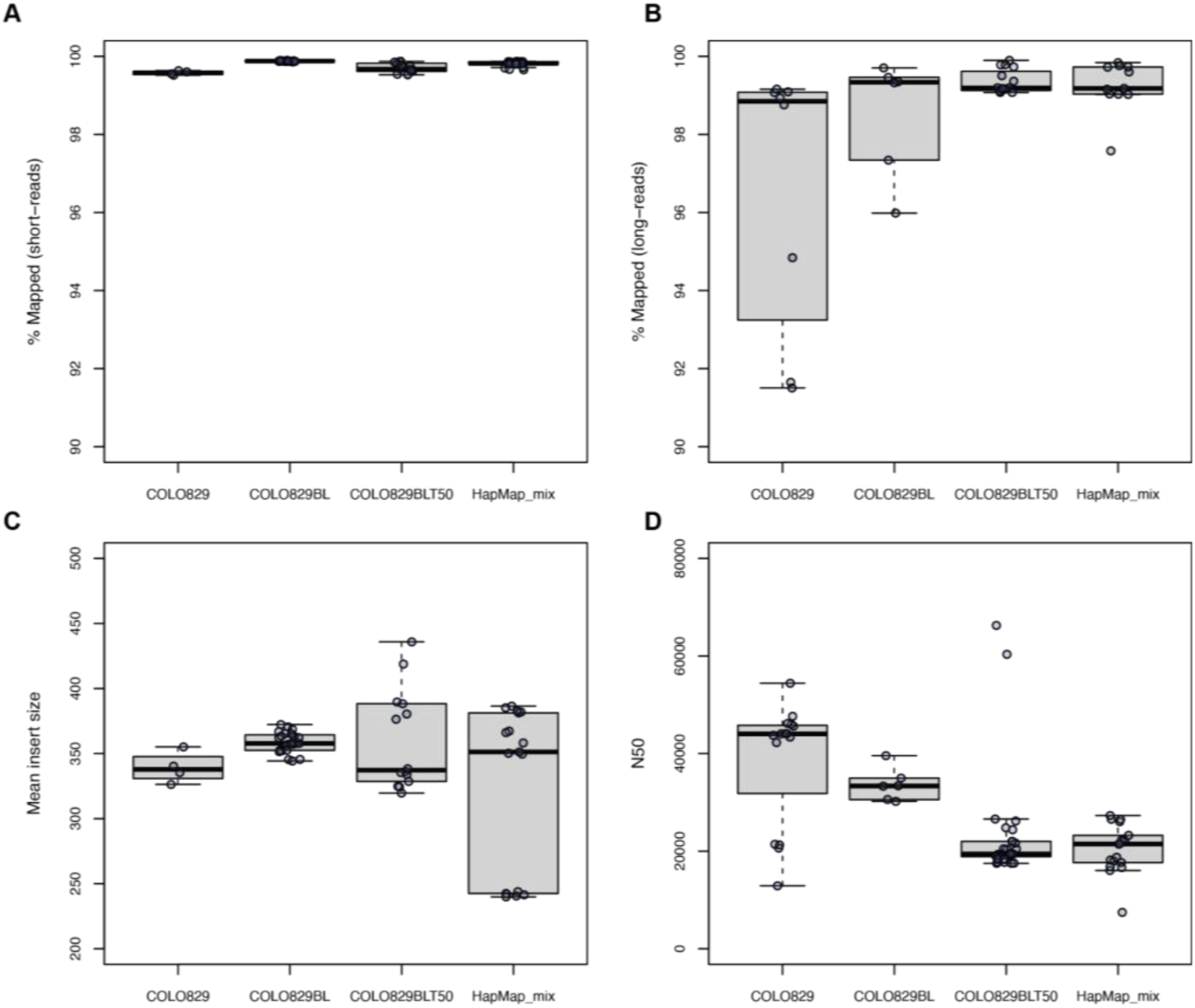
Overall mapping rates of Illumina short-read. (A) and PacBio, standard ONT long-read (B) bulk WGS data, as well as estimated insert sizes (C; for short-read WGS) and N50 (D; for long-read WGS) across the benchmark cell line samples analyzed in this study.

**Figure S2.**
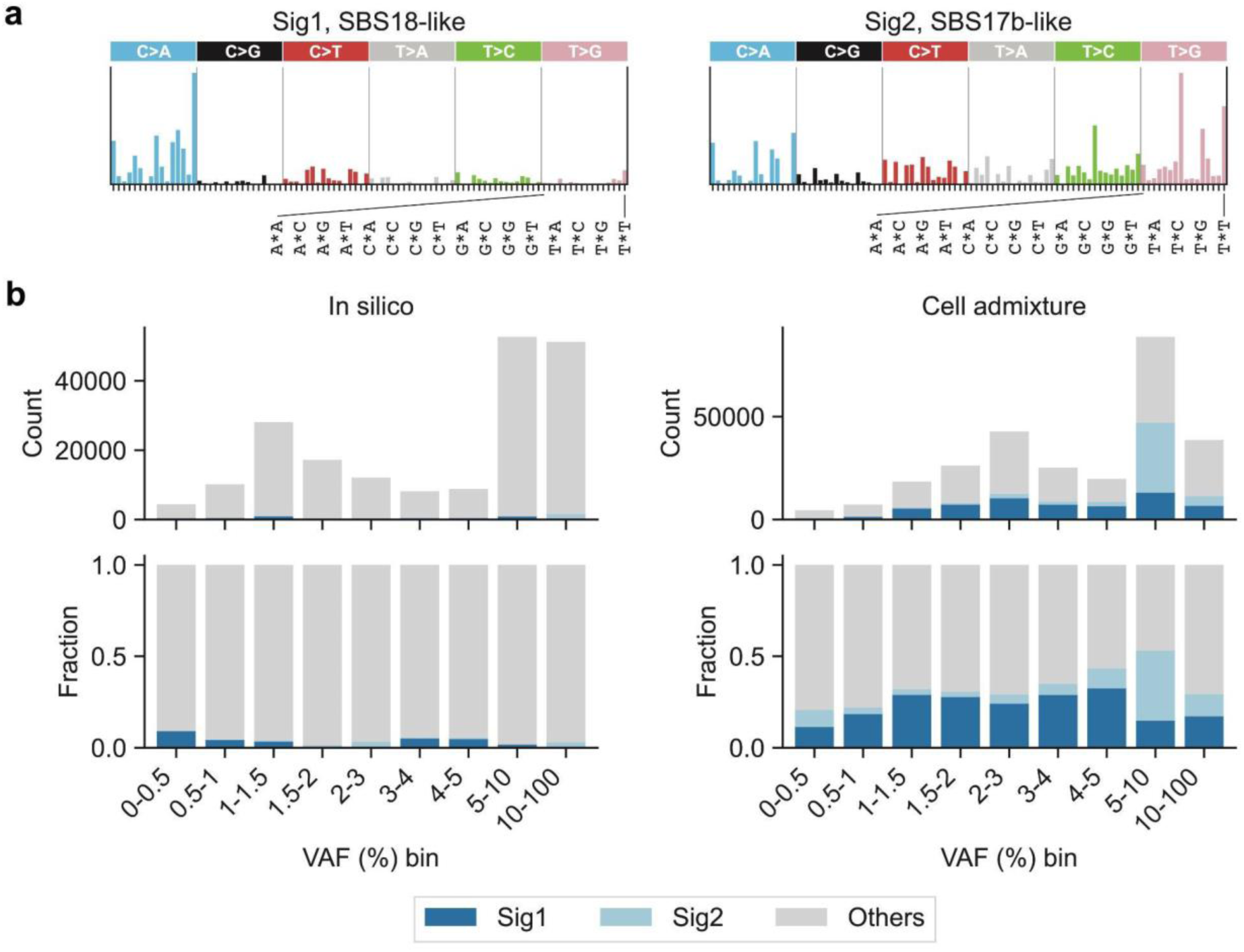
ROS-associated mutational signatures in cell admixture and in silico COLO829BLT50 datasets. **(a)** Two *de novo* mutational signatures likely associated with ROS damages in cell cultures. Signatures were *de novo* extracted from “nonvariant” false positive SNV calls stratified by sample, VAF bin, and variant calling pipeline. Sig1 (left) is similar to COSMIC SBS18 (cosine similarity = 0.93). Sig2 (right) shows strong peaks at N[T>G]T, especially C[T>G]T, which are characteristics of COSMIC SBS17b. Both SBS18 and SBS17b are annotated as ROS-associated in COSMIC. **(b)** Absolute (top) and relative (bottom) exposures of ROS-associated *de novo* mutational signatures in *in silico* (left) and cell admixture (right) COLO829BLT50 data. Exposures were calculated for “nonvariant” false positive SNV calls pooled over samples and variant calling pipelines.

**Figure S3.**
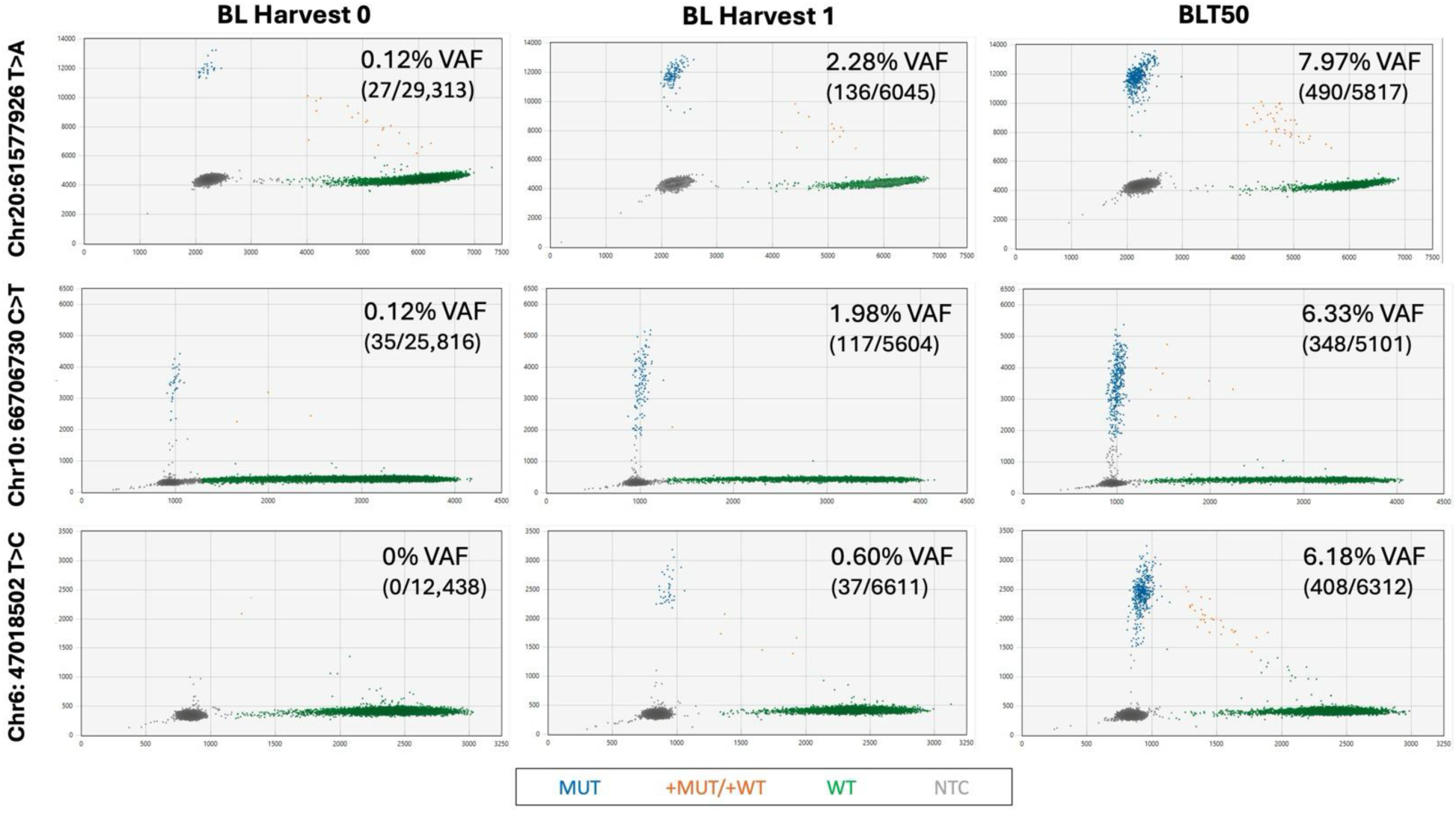
**Digital Droplet PCR (ddPCR) adjudication of variants found in the cell-culture derived set**. Three variants were selected for ddPCR sequencing. Variant allele frequencies for each sample are displayed in the upper right corner of each panel, along with the droplet counts below. Blue colored droplets indicate the homozygous presence of the variant, and orange colored droplets indicate a heterozygous presence, while green colored droplets show only WT allele at the position. Grey indicates drop out.

**Figure S4.**
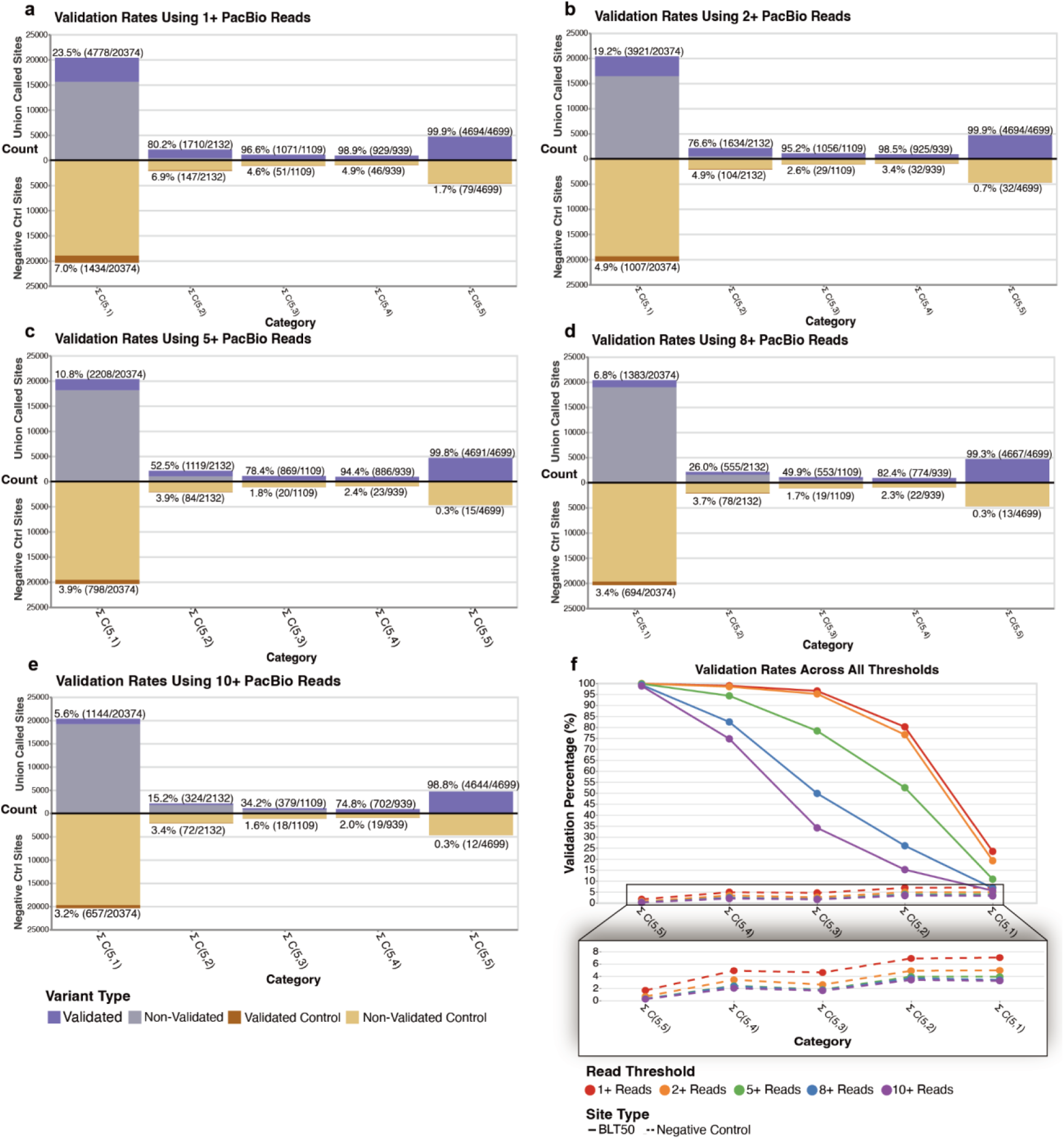
Validation rates for COLO829BLT50 admixture-only variants with increasing long-read coverage thresholds. For each histogram in panels (a-e), the top purple bars denote the counts of candidate and validated admixture-only variants. The bottom tan bars represent the same counts of candidate and validated alleles found in a negative control homogenate tissue which was sequenced using identical preparation methods and technology. Each histogram column represents the variants found in (**a**) single replicate (∑ C(5,1)), (**b**) two replicates (∑ C(5,2)), (**c**) three replicates (∑ C(5,3)), (**d**) four replicates (∑ C(5,4)), and (**d**) all five replicates (∑ C(5,5)). (**f**) comparison of the validation rates across increasing long-read thresholds, demonstrating that at our selected threshold of 2+, we maximize the amount of admixture-only variants validated, while minimizing the false positives that are also validated in the negative control.

**Figure S5.**
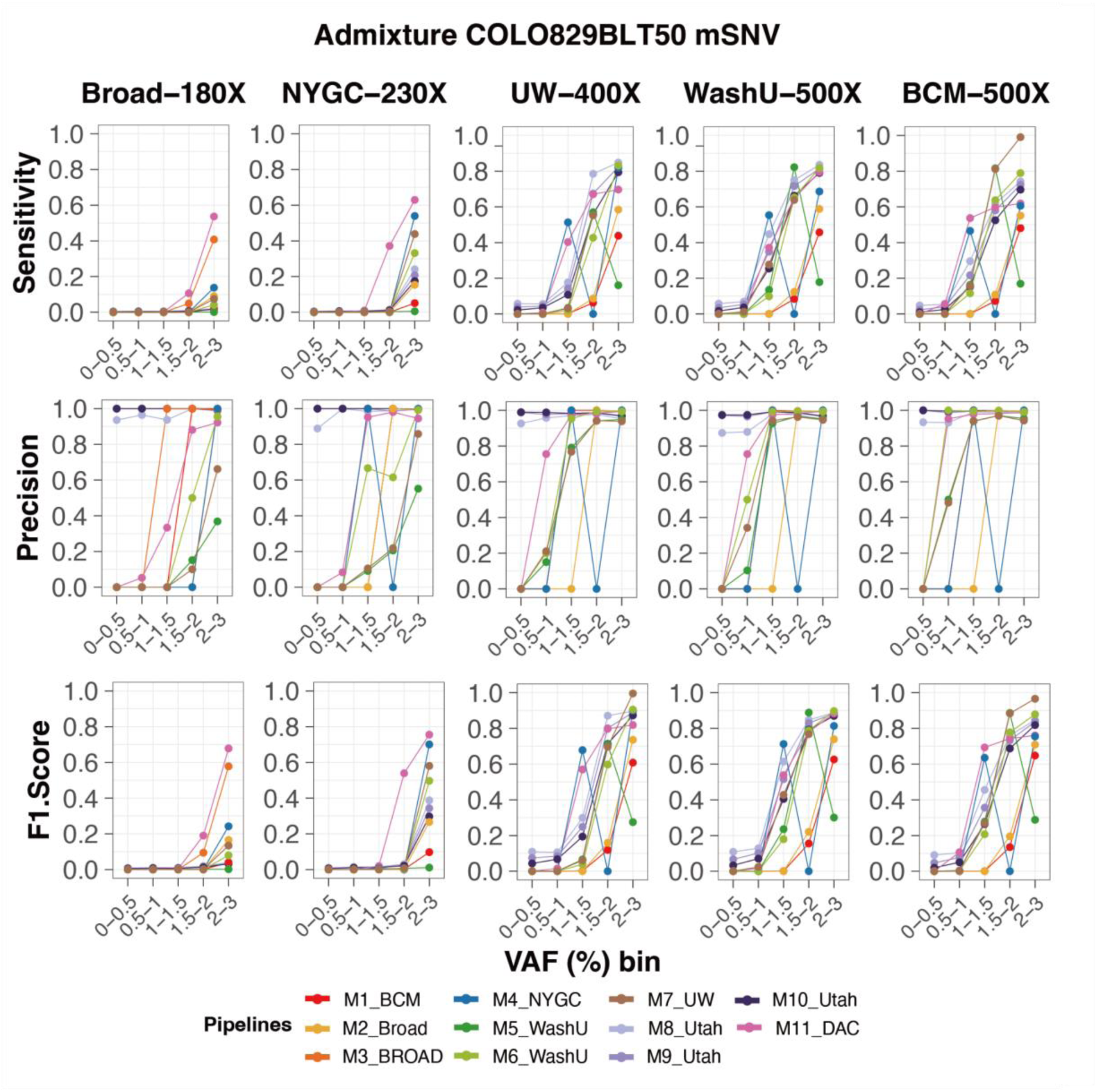
Performance evaluation of the mosaic SNVs with cell admixture COLO829BLT50. Sensitivity, Precision, and F1 score are shown with five independently sequenced COLO829BLT50. Ten or eleven detection pipelines were applied (**Table S3**) for mosaic SNVs in VAF below 3%.

**Figure S6.**
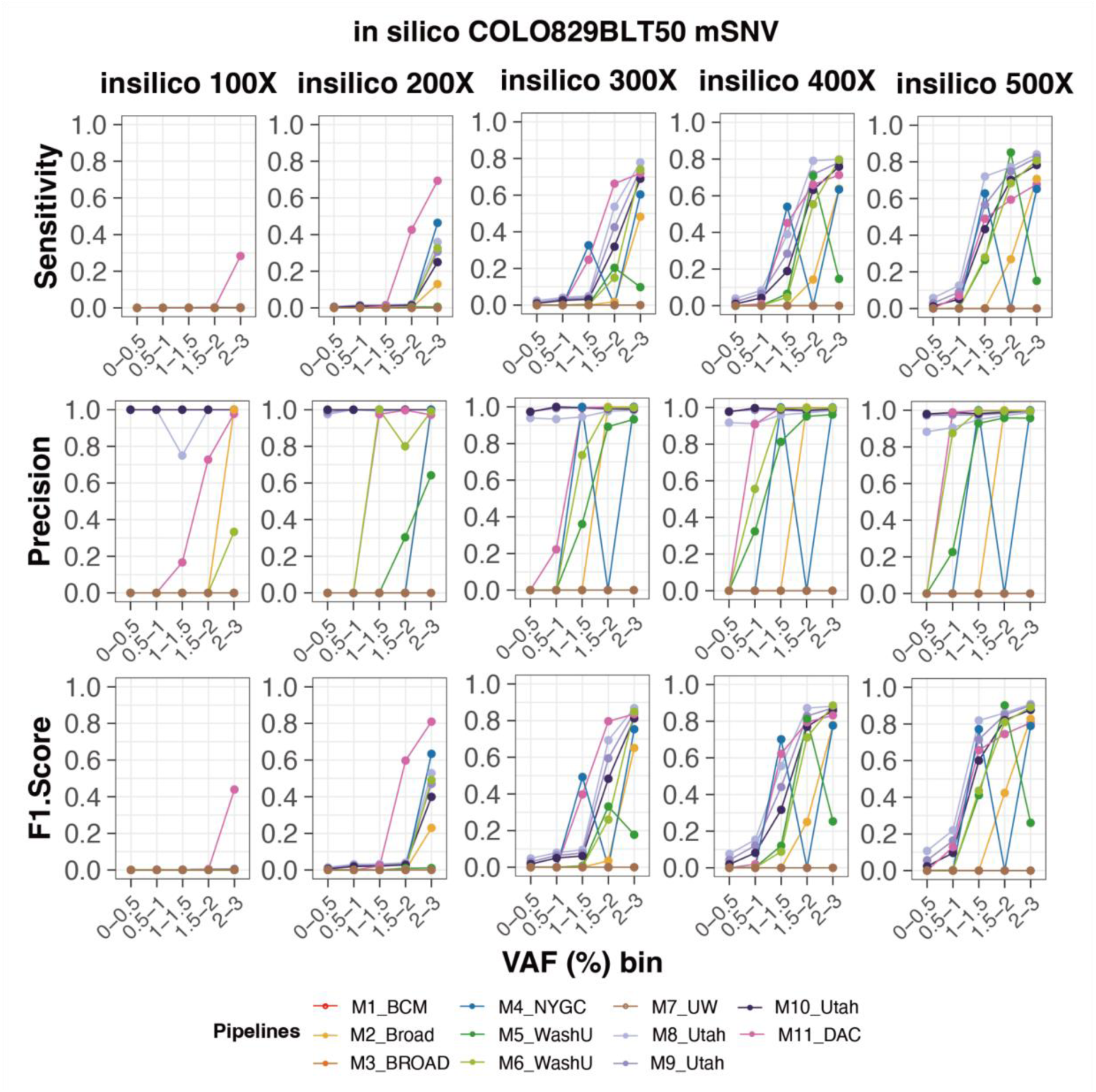
Performance evaluation of the mosaic SNVs with *in silico* COLO829BLT50. Sensitivity, Precision, and F1 score are shown with computationally generated COLO829BLT50 in five sequencing depths, ranging from 100× to 500×. Nine detection pipelines were applied (**Table S3**) for mosaic SNVs in VAF below 3%.

**Figure S7.**
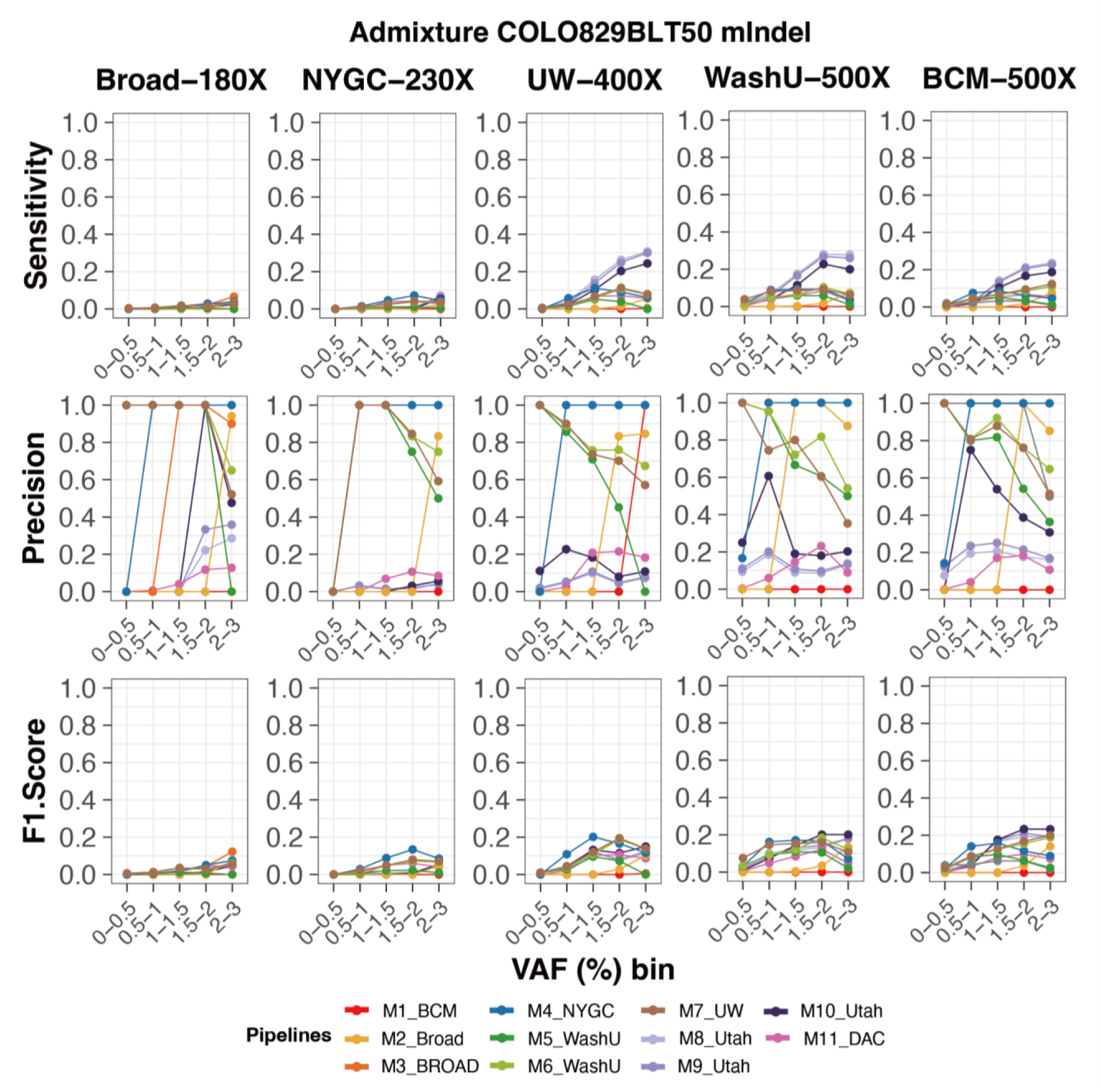
Performance evaluation of the mosaic Indels with cell admixture COLO829BLT50. Sensitivity, Precision, and F1 score are shown with five independently sequenced COLO829BLT50. Ten or eleven detection pipelines were applied (**Table S3**) for mosaic Indels in VAF below 3%.

**Figure S8.**
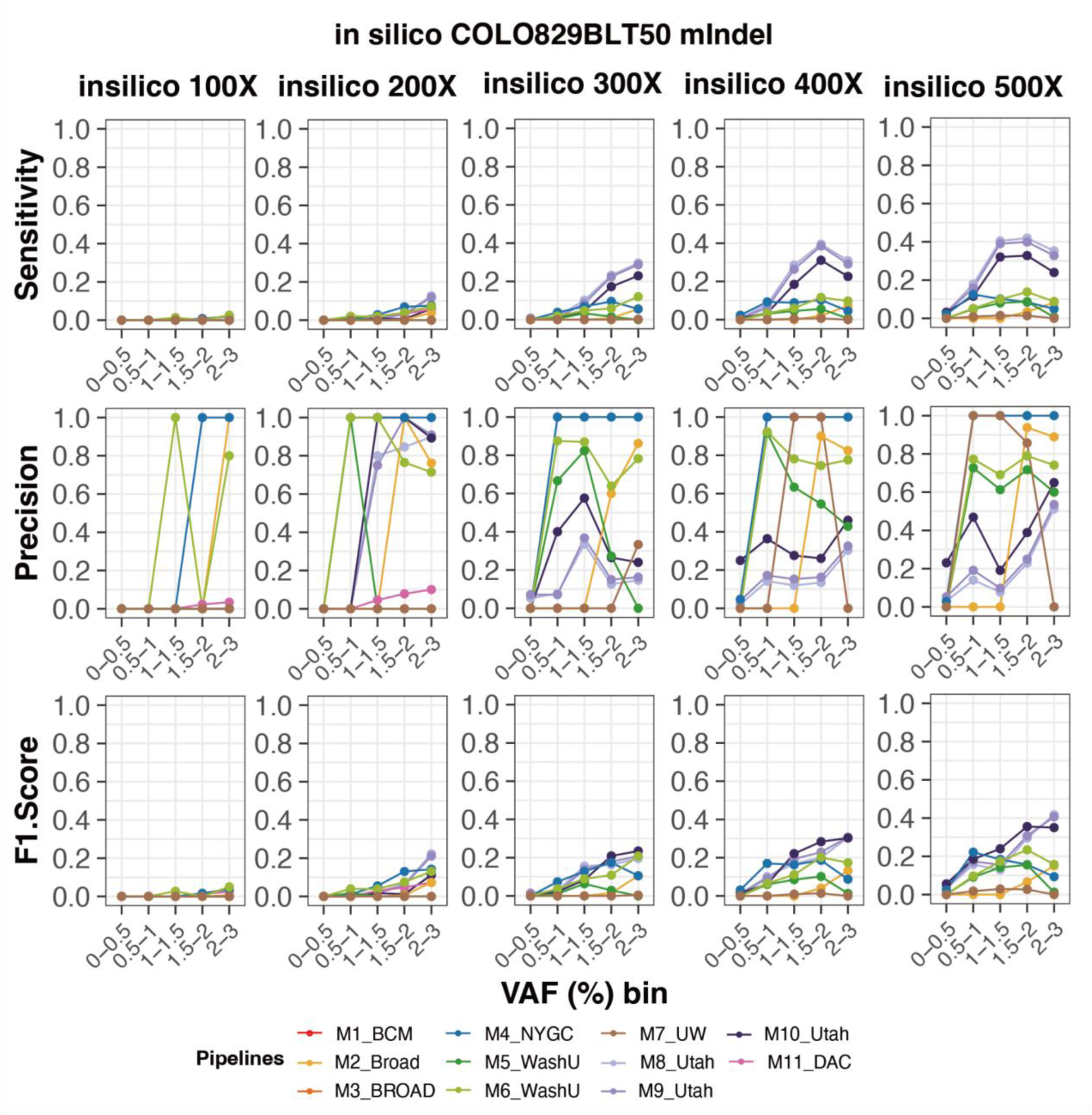
Performance evaluation of the mosaic Indels with *in silico* COLO829BLT50. Sensitivity, Precision, and F1 score are shown with five independently sequenced COLO829BLT50. Nine detection pipelines were applied (**Table S3**) for mosaic SNVs in VAF below 3%.

**Figure S9.**
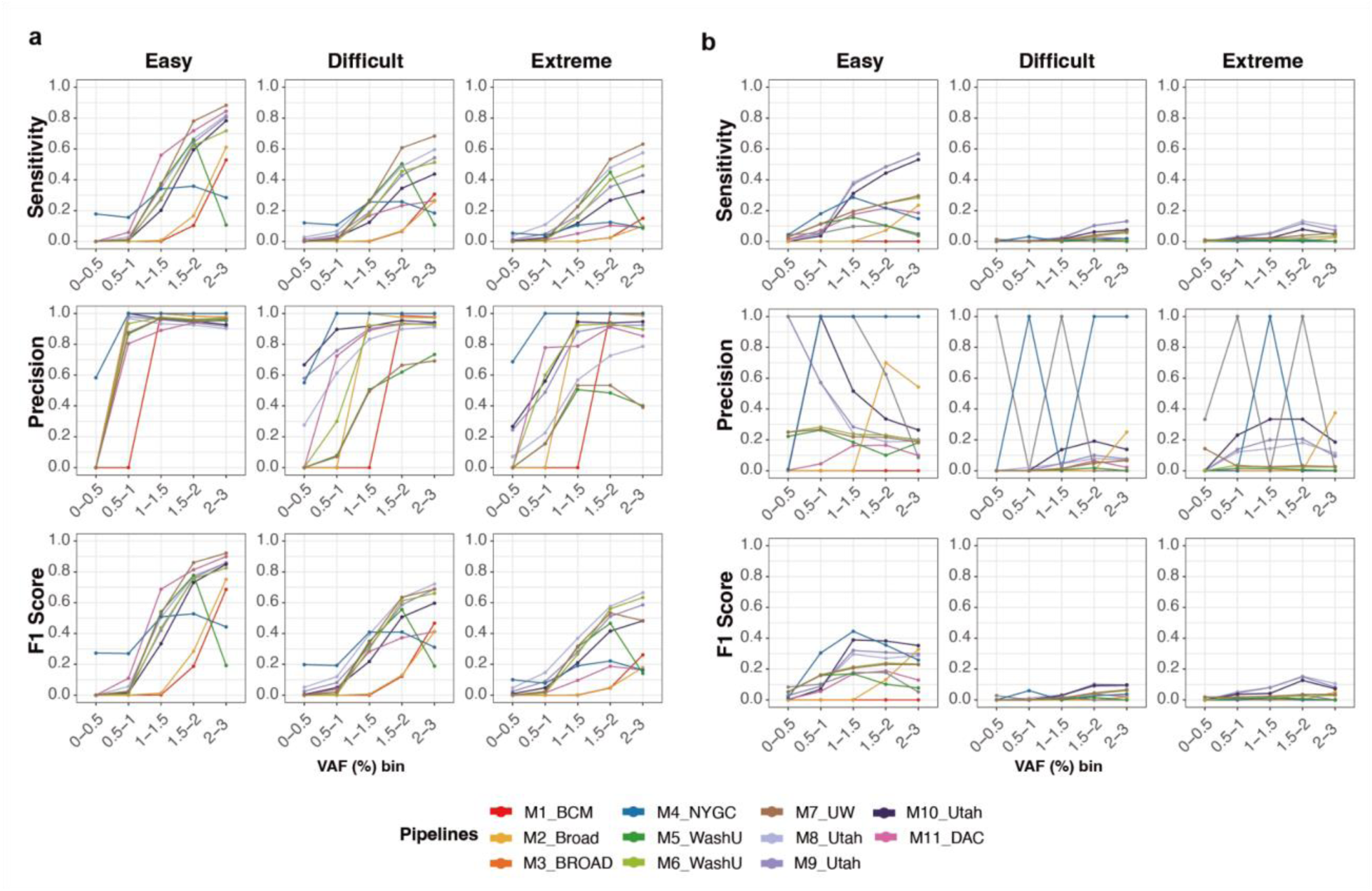
Performance evaluation of the mosaic SNVs and Indels in different genomic contexts. Sensitivity, Precision, and F1 scores are shown for ultra low-VAF (a) mSNVs and (b) mIndels. COLO829BLT50 with 500× coverage data was used, generated by BCM. Ten different pipelines were evaluated across easy, difficult, and extreme regions.

**Figure S10.**
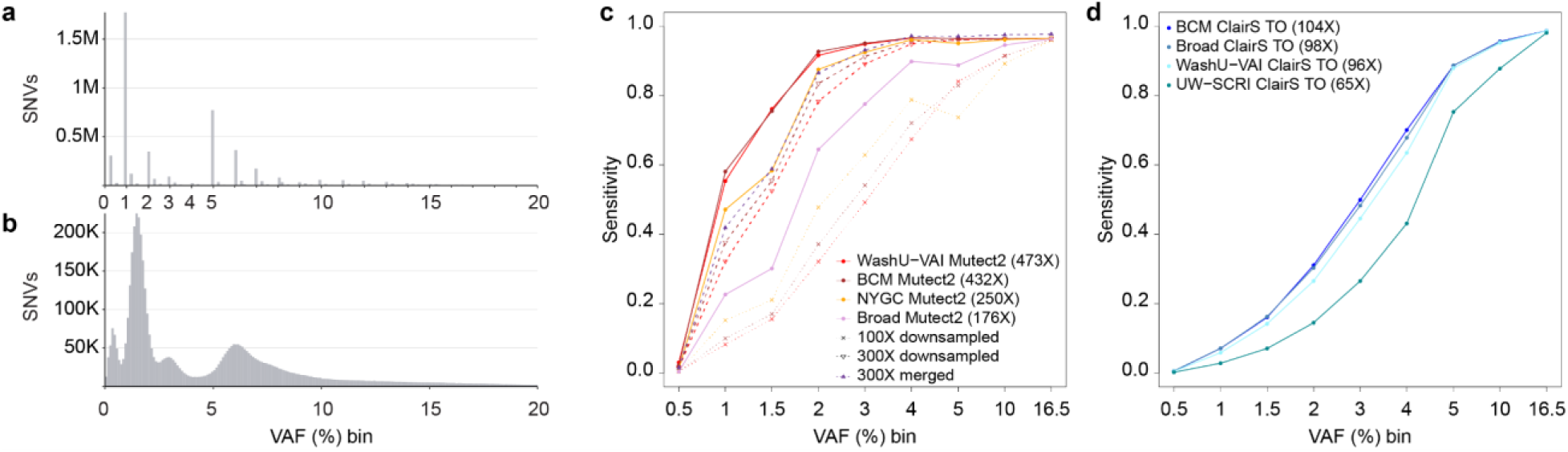
Expected and observed VAF distribution and detailed sensitivity in HapMap samples. Distribution of expected VAFs (a) and observed VAFs (b) in the HapMap mixture, as calculated by pileup at target positions. (c) Sensitivity of somatic variant calling with Mutect2 across expected VAF bins in Illumina short-read HapMap samples, including datasets downsampled to 300× and 100× coverage, as well as merged to 300×.(d) Sensitivity of somatic variant calling with ClairS TO across expected VAF bins per PacBio long-read sample.

## Supplementary tables

**Table S1:** Overview of sequencing depth of short-read and long-read datasets used for analysis of COLO289 and HapMap mixture benchmarking experiments. Sample Center llumina short-read WGSPacBio long-read WGS.

| Sample | Center | Illumina short-read WGS | PacBio long-read WGS |
| --- | --- | --- | --- |
| COLO829 | UW-SCRI-GCC | 367.9X | 168.4X |
|  | TOTAL | 367.9X | 168.4X |
| COLO829BL | UW-SCRI-GCC | 277.9X | 325.5X |
|  | TOTAL | 277.9X | 325.5X |
| COLO829BLT50 | BCM-GCC | 474.9X | 106.4X |
|  | Broad-GCC | 166.6X | 156.5X |
|  | NYGC-GCC | 213.6X | -- |
|  | UW-SCRI-GCC | 385.4X | 23.8X |
|  | WashU-VAI-GCC | 471.5X | 102.7X |
|  | TOTAL | 1,712.0X | 389.5X |
| HapMap mixture | BCM-GCC | 432.5X | 103.6X |
|  | Broad-GCC | 167.2X | 98.1X |
|  | NYGC-GCC | 250.0X | -- |
|  | UW-SCRI-GCC | -- | 65.4X |
|  | WashU-VAI-GCC | 473.1X | 95.6X |
|  | TOTAL | 1,322.8X | 362.7X |

**Table S2:** PacBio validation rate of short-read-based SNV and Indel calls in COLO829BL for different variant calling tools.

| Method | Mutation type | Total Illumina call | Alt in COLO829BL | No alt in COLO829 | Valid | Valid rate (%) | No alt in COLO829 rate (%) | Alt in COLO829BL rate (%) |
| --- | --- | --- | --- | --- | --- | --- | --- | --- |
| **Mutect2** | SNV | 44,623 | 1,349 | 624 | 42,476 | 95.19 | 1.40 | 3.02 |
| **Strelka2** |  | 55,673 | 5,608 | 5,642 | 43,716 | 78.52 | 10.13 | 10.07 |
| **RUFUS** |  | 38,790 | 993 | 380 | 37,316 | 96.20 | 0.98 | 2.56 |
| **VarNet** |  | 44,288 | 1,114 | 3,208 | 39,693 | 89.62 | 7.24 | 2.52 |
| **Mutect2** | Indel | 2,416 | 35 | 650 | 1,675 | 69.33 | 26.90 | 1.45 |
| **Strelka2** |  | 2,095 | 134 | 767 | 1,194 | 56.99 | 36.61 | 6.40 |
| **RUFUS** |  | 2,099 | 355 | 714 | 1,008 | 48.02 | 34.02 | 16.91 |
| **VarNet** |  | 800 | 6 | 65 | 729 | 91.13 | 8.13 | 0.75 |

**Table S3:**
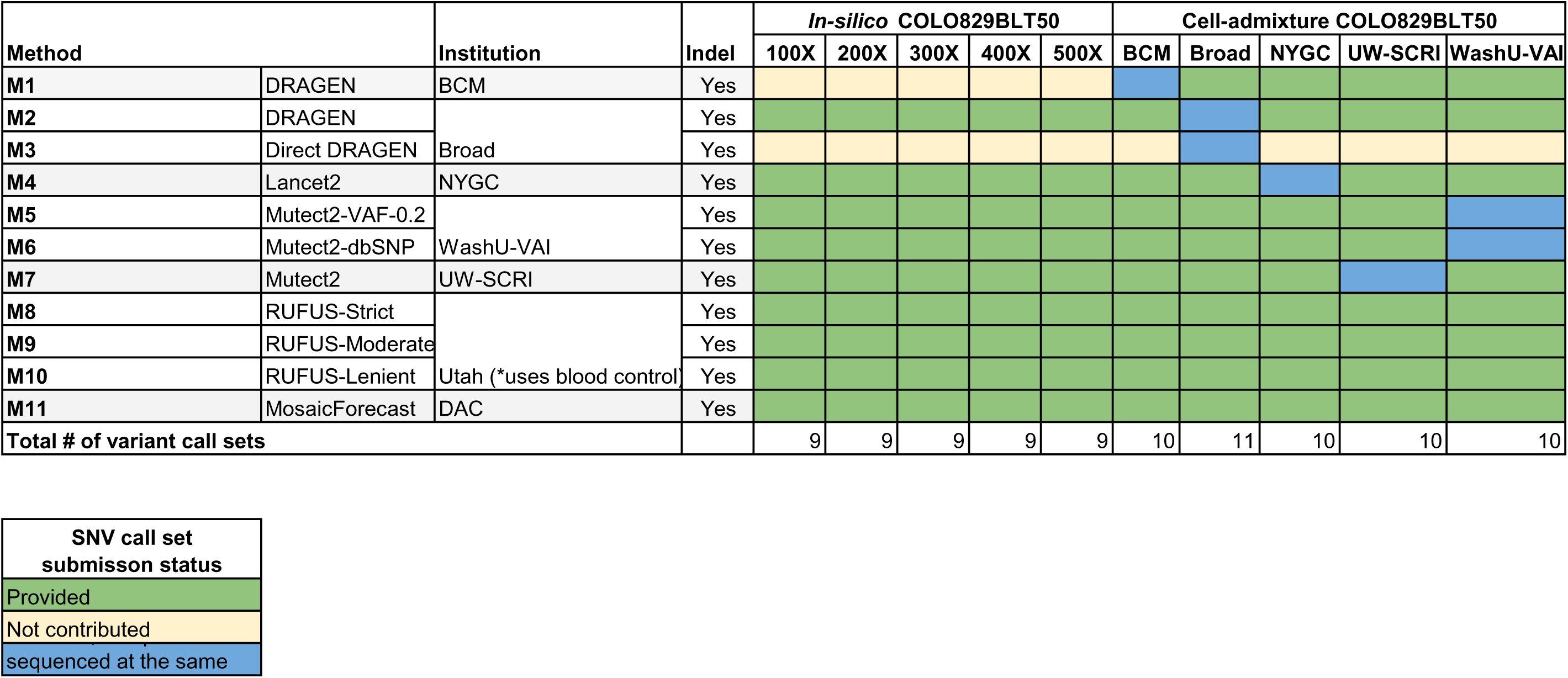
Variant calling methods applied to *in-silico* and cell-admixture COLO829BLT50 samples by participating institutions.

**Table S4:** F1 Score average across different VAF ranges and sequencing depths.

|  | Sequencing Depth |  |  |  |  |
| --- | --- | --- | --- | --- | --- |
|  | 100x | 200x | 300x | 400x | 500x |
| **VAF < 1%** | 0 | 0 | 0.00024531 | 0.00967606 | 0.06414392 |
| **1% < VAF < 3%** | 0.14807572 | 0.476122 | 0.67649267 | 0.74948148 | 0.73641945 |
| **gain VAF < 1%** | NA | 0 | 0.00024531 | 0.00943075 | 0.05446786 |
| **gain 1% < VAF < 3%** | NA | 0.32804628 | 0.20037068 | 0.07298881 | -0.013062 |
| **Gain percentage in VAF < 1%** | NA | NA | NA | 38.44421344 | 5.629136239 |
| **Gain percentage in 1% < VAF < 3%** | NA | 2.215395475 | 0.420838945 | 0.10789298 | -0.017428049 |

**Table S5:** Population code and number of germline SNVs within confident regions per individual HapMap cell line contributing to the HapMap mixture.

| HapMap cell line | Population code | Number of SNVs |
| --- | --- | --- |
| HG002 | EUR/ASH JEW | 3,176,093 |
| HG005 | CHB | 3,143,272 |
| HG00438 | EAS | 3,168,150 |
| HG02257 | AFR | 3,816,806 |
| HG02486 | AFR | 3,843,750 |
| HG02622 | AFR | 3,886,058 |

